# Circulating mucosal-like IgA responses associate with severity of Puumala orthohantavirus-caused hemorrhagic fever with renal syndrome

**DOI:** 10.1101/2024.07.30.605778

**Authors:** Luz E. Cabrera, Cienna Buckner, Veronica Martinez, Sanna Mäki, Olli Vapalahti, Antti Vaheri, Jussi Hepojoki, Johanna Tietäväinen, Satu Mäkelä, Jukka Mustonen, Tomas Strandin

## Abstract

Old World Orthohantaviruses cause hemorhagic fever with renal syndrome (HFRS) characterized by increased vascular permeability and acute kidney injury (AKI). Despite the systemic nature of the disease, the virus enters humans through inhalation and therefore initially encounters the immunoglobulin class A (IgA) dominated mucosal immune system. Herein, we characterized systemic IgA responses and their potential relationship to the mucosal immune activation by examining blood samples obtained from patients hospitalized due to acute Puumala orthohantavirus infection. Our findings reveal increased frequencies of IgA-expressing circulating mucosal-associated B1 cells and plasmablasts, as well as elevated levels of polyreactive, polymeric, virus-specific and secretory IgA in the acute stage of the disease. Importantly, the levels of circulating virus-specific and secretory IgA associated with the severity of AKI. Furthermore, circulating polymeric IgA displayed enhanced effector functions by forming stable complexes with the IgA receptor CD89 and induced pro-inflammatory neutrophil responses. These results suggest that, while an efficient mucosal immune response is likely to be crucial for infection clearance, an excessive mucosal immune activation may contribute to HFRS disease progression.

## Introduction

Orthohantaviruses endemic in Europe and Asia cause hemorhagic fever with renal syndome (HFRS), whereas the viruses present in the Americas cause hantavirus cardiopulmonary syndrome (HCPS) (Vial, Ferrés et al. 2023). Puumala orthohantavirus (PUUV) is the most prevalent orthohantavirus in Europe and carried by the bank vole (Myodes glareolus). When transmitted from rodent excretions to humans, it causes a relatively mild form of HFRS (PUUV-HFRS), the typical symptoms of which include fever, vomiting, diarrhea, headache, backache and myopia (Vaheri, Smura et al. 2023). Hospitalized PUUV-HFRS patients often present with acute kidney injury (AKI), the molecular mechanisms of which remain unclear. Increased vascular permeability due to the loss of endothelial cell (EC) barrier functions is pathognomonic to orthohantavirus-mediated diseases, including PUUV-HFRS. Virus replication occurs primarily in the capillary ECs but is without direct cytopathic effects and thus does not directly explain the increased vascular permeability. In fact, it is generally believed that aberrant immunological responses towards the virus play a major role in the development of HFRS (Klingström, Smed-Sörensen et al. 2019). However, the lack of an animal disease model greatly hampers the detailed pathogenesis investigations for HFRS.

The virus infection occurs through inhalation, which suggests that the mucosal immune system of the airways is the first to respond to the infection. Furthermore, HFRS symptoms include vomiting and diarrhea, which implies the involvement of the gastrointestinal tract that harbors the largest mucosal immune compartment in humans. Mucosal immunity is largely mediated by IgA, the most abundant antibody isotype in humans. A major proportion of IgA is produced by mucosal tissue resident B cells in the lamina propria, from where it migrates through transepithelial transport to the luminal space of the mucosal tissue (Sinha, Yaugel-Novoa et al. 2024). During the transport, the secretory component (SC) is covalently linked to IgA, allowing the distinction of secretory IgA (sIgA) from the other IgA forms. The mucosal IgA is distinct from systemic circulating IgA due to its propensity to form dimers or polymers (Matsumoto 2022) and its ability to bind multiple antigens simultaneously (i.e. polyreactivity) (Bunker, Erickson et al. 2017).

The trafficking of antibody secreting cells (ASCs) to the intestine is mediated by their expression of the homing receptors α4β7 integrin (Kunkel and Butcher 2003) and C-C Chemokine receptor type 9 (CCR9) (Hieshima, Kawasaki et al. 2004), with the latter being responsible for homing specifically to the small intestine. The fast-responding and rigorously IgA expressing intestinal plasma cells are further characterized by their expression of integrin CD11b (Kunisawa, Gohda et al. 2013). In addition to plasmablasts (PBs) and plasma cells, B1 cells are another class of fast-responding ASCs (Suchanek and Clatworthy 2023). Originally discovered in the peritoneal cavity of mice, B1 cells are innate-like cells that spontaneously release antibodies (abs) with minimal somatic hypermutation, producing natural abs distinct from the high-affinity abs produced by plasma cells. While the exact phenotype of human B1 cells has been a topic of debate (Sanz, Wei et al. 2019), Quach *et al* were able to reliably discriminate B1 cells from other ASCs as CD20^+^CD27^+^CD38^-/int^CD43^+^CD70^-^ (Quách, Rodríguez-Zhurbenko et al. 2016). Interestingly, in response to infection, B1 cells migrate to the lymph nodes through increased expression of CD11b (Waffarn, Hastey et al. 2015).

We recently observed increased frequencies of IgA-expressing PBs in the circulation of acute PUUV-HFRS (Hepojoki, Cabrera et al. 2021). However, the role of IgA and mucosal immunity in the development of PUUV-HFRS is not well understood. In the current study, we aimed to gain a deeper view of the IgA-dominated mucosal immunity activation in HFRS by analyzing mucosal homing receptor expression of IgA ASCs and assessing mucosal-associated IgA levels in the circulation during acute, convalescent and recovery stage PUUV-HFRS. Furthermore, we studied the potential pathologic and pro-inflammatory roles of circulating IgA during the disease.

## Materials and methods

### Patient population

The study material consisted of serum and PBMCs from serologically-confirmed PUUV-infected patients hospitalized (Tampere University Hospital, Finland) between 2005 and 2009. The patient samples were systematically collected from a cohort of 55 individuals (or 25 in case of PBMCs), at different time points, which included the acute phase for samples taken during hospitalization (5-9 days after onset of fever {aof}), the postacute/convalescent phase for samples taken 14 days after discharge from hospital (∼30 days aof), and the recovery phase for samples taken 6 months and one year aof. The recovery samples served as steady state controls. The study protocol was approved by the Ethical Committee of Tampere University Hospital with permit number R04180.

Standard methods were employed to determine daily white blood cell (WBC) and thrombocyte count, plasma C-reactive protein (CRP), and serum creatinine concentrations at the Laboratory Centre of the Pirkanmaa Hospital District (Tampere, Finland). Descriptive patient characteristics are given in Table1.

### PBMC analysis by flow cytometry

Frozen PUUV patient PBMCs were thawed in a 37°C water bath for 5 min and washed with R10 (RPMI-1640 (Sigma Aldrich) supplemented with 10% inactivated FCS (Gibco), 100 IU/ml of penicillin (Sigma Aldrich), 100 μg/ml of streptomycin (Sigma Aldrich) and 2 mM of L-glutamine (Sigma Aldrich)) and including 100 μg/ml DNAse I (Sigma Aldrich). After a 10 min incubation at RT, the cells were washed once with the conditioned medium and once with PBS-EDTA (PBS with 2 mM EDTA), followed by cell quantification using Bio-Rad cell counter TC20.

One to three million PBMCs were incubated in 1% FCS and FcR blocking reagent (BioLegend), and the cells stained for 30 min at RT with a cocktail of fluorescent dye conjugated anti-human mouse monoclonal antibodies recognizing cell surface antigens and a dead cell marker (the antibody panel is provided in detail in Table 2). After staining, the cells were washed once with PBS-EDTA and fixed with 1% paraformaldehyde before FACS analysis with a 4-laser (405, 488, 561 and 637 nm) Quanteon Novocyte (Agilent Technologies).

**Table 1.** Clinical and laboratory data during hospitalization in patients with acute PUUV infection.

| Parameter (unit) | Range (Median± IQR)<br>Serum (n = 55) | Range (Median + IQR)<br>PBMCs (n = 25) | Reference value |
| --- | --- | --- | --- |
| Age (years) | 22 – 74 (39 ± 19) | 25 – 67 (35 ± 22) | – |
| Gender (Male:Female) | 33:22 | 15:10 | – |
| Duration of fever (days) | 3 – 15 (8 ± 3) | 3 – 9 (7 ± 2) | – |
| Length of hospital stay<br>(days) | 2 – 25 (5 ± 3) | 2 – 25 (5 ± 2) | – |
| Max Creatinine (μmol/L) | 51 – 1499 (165 ± 376) | 51 – 1499 (150 ± 279) | Male: 62 – 115 /<br>Female: 53 – 97 |
| Max WBC (E9/L) | 6 – 26 (10 ± 5) | 7 – 26 (11 ± 5) | 3.4 – 8.2 |
| Min Thrombocytes (E9/L) | 15 – 118 (54 ± 41) | 15 – 118 (61 ± 44) | 150 – 360 |
| Max C-reactive protein<br>(mg/L) | 16 – 267 (91 ± 67) | 16 – 267 (89 ± 92) | < 4 |

WBC = white blood cells, IQR = Interquartile range

**Table 2.** PBMC flow cytometry antibody panel.

| Marker | Fluorochrome | Clone | Company |
| --- | --- | --- | --- |
| CD3 | FITC | OKT3 | ThermoFisher |
| CD14 | FITC | MEM-18 | Immunotools |
| CD56 | FITC | MEM-188 | Immunotools |
| CD66 | FITC | 6g5j | Immunotools |
| $\alpha 4\beta 7$ | PE | Hu117 | RnD Systems |
| IgA | PE-Cy7 | IS11-8E10 | MACS Miltenyi |
| CD43 | PerCP-eFluor 710 | 4-29-5-10-21 | ThermoFisher |
| CCR9 | APC | L053E8 | Biolegend |
| IgD | APC-Cy7 | IA6-2 | Biolegend |
| **CD38** | BV421 | HB7 | Biolegend |
| **CD19** | BV510 | SJ25C1 | Biolegend |
| **CD11b** | BV605 | ICRF44 | BD Biosciences |
| **CD27** | BV711 | M-T271 | Biolegend |
| **CD70** | BV786 | Ki-24 | BD Biosciences |
| **CD20** | BB615 | 2H7 | Bio-Rad |

#### Neutrophil analysis by flow cytometry

Neutrophils were isolated from EDTA-anticoagulated blood of healthy controls using Polymorphprep (Proteogenix) and 2*10^6^/ml neutrophils in R10 were incubated with 10 µg/ml IgA in 10 µl for 4 hours at 37 °C. Samples were then incubated with CellROX green dye (Thermo Fisher) and an antibody cocktail (Table 3) for 30min at RT. Finally, the samples were diluted 1:20 in PBS and analyzed using a BD Fortessa LSRII flow cytometry (BD Biosciences).

**Table 3.** Neutrophil flow cytometry antibody panel.

| Marker | Fluorochrome | Clone | Company |
| --- | --- | --- | --- |
| **LOX-1** | PE | 15C4 | BioLegend |
| **IgA** | PE-Cy7 | IS11-8E10 | MACS Miltenyi |
| **CD11b** | APC-Cy7 | ICRF44 | Biolegend |
| **CD66b** | BV421 | G10F5 | BD Biosciences |
| **CD62L** | BV605 | DREG-56 | BD Biosciences |
| **HLA-DR** | BV786 | G46-6 | BD Biosciences |

#### PBMC cultures

Frozen patient PBMCs were revived similarly as described for flow cytometry above. PBMCs (at 1 million/ml) were cultured in R10 for 6 days at 37 °C, centrifuged for 10 min at 400 g and supernatants collected for ELISA as specified below.

#### ELISAs

The amounts of IgA and sIgA were measured by ELISA protocols including capture mAbs anti-human IgA (clone MT57, Mabtech) for IgA and anti-human secretory component (clone SC-05, Exbio) for sIgA. Capture mAbs were coated onto MaxiSorp plates (ThermoFisher) at 2 µg/ml in PBS overnight in +4°C and blocked with 1 % BSA-PBS for 1 hr prior to incubating with diluted patient blood samples for 1 hr. After washing with PBS containing 0.05% Tween-20 (PBST), the plates were incubated with Horseradish peroxidase (HRP, Dako, dilution 1:6000)- or alkaline phosphatase (ALP, clone MT20, Mabtech, dilution 1:1000)-conjugated secondary IgA-specific antibodies for IgA and sIgA. After washing with PBST, the HRP- and ALP-conjugated Abs were detected by 3,3′,5,5′-tetramentylbenzidine (TMB) or para-nitrophenyl phosphate (pNPP) substrates, respectively (Thermo Scientific). TMB-mediated reaction was stopped by addition of 0.5 M H_2_SO_4_. Absorbances (450 nM for HRP- and 405 nM for ALP-based reactions) were read by HIDEX sense microplate reader. As standards, native human IgA from Mabtech and sIgA isolated from healthy donor saliva were used. To detect virus-specific, polyreactive or receptor binding of IgA, the capture antibodies were replaced by baculovirus-expressed PUUV nucleocapsid (N) protein (2 µg/ml), DNP-conjugated albumin (10 µg/ml, Sigma) or CD89 (2 µg/ml, RnD systems), respectively.

#### IgA isolations

IgA was isolated from serum using peptide M agarose following manufacturer’s recommendations (Invivogen). The isolated IgAs were eluted into 10 mM Glycine pH 1.7, followed by buffer exchange to PBS and sample concentration using 50 kDa centrifugal filters (Amicon, Millipore). Approximately 1 mg of isolated IgAs were separated into polymeric, dimeric and monomeric fractions by size-exclusion chromatography (Sephadex G-200) using Äkta HPLC (Cytiva), fractions concentrated using 50 kDa centrifugal filters and analyzed by ELISA.

#### Binding kinetics

The binding of IgA to CD89 was analyzed with surface plasmon resonance with Biacore T100 instrumentation (Cytiva). The flow cells of a CM5 S series biosensor chip (Cytiva) were covalently coated with CD89 via standard amine coupling or left empty (negative control). The coating with CD89 resulted in a resonance unit (RU) increase of approximately 1000. The binding was analyzed using PBS as the running buffer and a flow rate of 20 μL min–1. The kinetics of the IgA binding to CD89 were measured by varying IgA concentrations with a contact time of 2 min and a dissociation phase of 10 min. After the completion of the dissociation phase, the flow cells were regenerated with 10 mM Glycine, pH 1.7. The data were evaluated by first subtracting the sensorgrams obtained from empty negative control flow cell from those obtained from the flow cells containing CD89. Association and dissociation rate constants were obtained by the Langmuir global fit model (BiaEvaluation Software, Cytiva).

#### On cell-ELISA

Blood microvascular endothelial cells (BECs) were obtained from Lonza and maintained in endothelial basal medium (EBM-2) supplemented with SingleQuots™ Kit containing 5% fetal bovine serum (FBS), human endothelial growth factor, hydrocortisone, vascular endothelial growth factor, human fibroblast growth factor-basic, ascorbic acid, R3-insulin like growth factor-1, gentamicin and amphotericin-B (Lonza). The experiments were performed on cells at passage 7 or 8. Isolated IgAs at 10 µg/ml were incubated with BECs for 45 min on ice, after which cells were washed with PBS 3 times and fixed with 4% paraformaldehyde for 10 min at RT. Fixed cells were blocked with 1% BSA-PBS for 30 min at RT and incubated with HRP-conjugated polyclonal anti-human IgA (1:6000, Dako) for 1 hr and washed with PBS 3 times. Color was developed with TMB and reaction stopped with 0.5 M H_2_SO_4._ Absorbance at 450nm was read by HIDEX sense microplate reader.

#### Statistical analyses

Statistically significant differences between time points were assessed by generalized estimating equations with working correlation matrix set as independent and gamma with log link scale response as the model (SPSS v29). Grouped non-parametric patient data was analyzed Mann-Whitney test and differences between treatment groups were analyzed by two-way ANOVA using Tukey’s test for multiple comparisons using GraphPad Prism (v8).

## Results

### IgA ASCs are significantly increased during acute PUUV-HFRS

We previously identified significantly increased frequencies of IgA-expressing PBs in the peripheral blood during acute PUUV-HFRS (Hepojoki, Cabrera et al. 2021). To understand the tissue tropism of these PBs, we developed an assay to measure the surface expression of mucosal homing receptors integrin α4β7, CD11b, and CCR9 in virus-induced PBs, using multicolor flow cytometry. Alongside PBs (CD20^±^CD19^+^IgD^-^CD27^+^CD38^++^CD43^+^), we investigated the frequencies and IgA expression of circulating mucosal-associated B1 cells (CD20^+^CD19^+^CD27^+^CD38^-/int^CD43^+^CD70^-^, representative gating strategy shown in Supplementary Fig. 1).

In agreement with our previous findings, we observed a significant increase (at least 10-fold) in PB frequencies during acute PUUV-HFRS as compared to recovery stage samples collected 360 days after fever onset (R360), which represented healthy steady state (Fig. 1A). Interestingly, a significantly diminished proportion of PBs expressed CCR9 during acute PUUV-HFRS as compared to steady state (Fig 1B), and the frequencies of α4β7 expressing PBs were also reduced at 6 days after onset of fever (aof) (Fig. 1C). In contrast, the proportion of CD11b-expressing PBs significantly increased during acute PUUV-HFRS (Fig. 1D). Of all the PBs detected during the acute stage approximately 20-30% expressed surface IgA, while steady state samples demonstrated over 30% surface IgA expression (Supplementary Fig 2A). We previously observed that ∼50 % of acute PUUV infection-associated PBs expressed intracellular IgA (Hepojoki, Cabrera et al. 2021), which together with the current findings indicated that a significant proportion of IgA PBs did not express surface-bound IgA. Compared to all PBs, the observed trends in the expression of α4β7 and CD11b in IgA PBs were comparable, showing diminished early α4β7 expression and significantly elevated CD11b expression in acute PUUV-HFRS (Supplementary Fig2. B-D). However, the frequencies of IgA-expressing PBs were too low for a robust comparison of differences in CCR9 expression between the time points.

**Figure 1.**
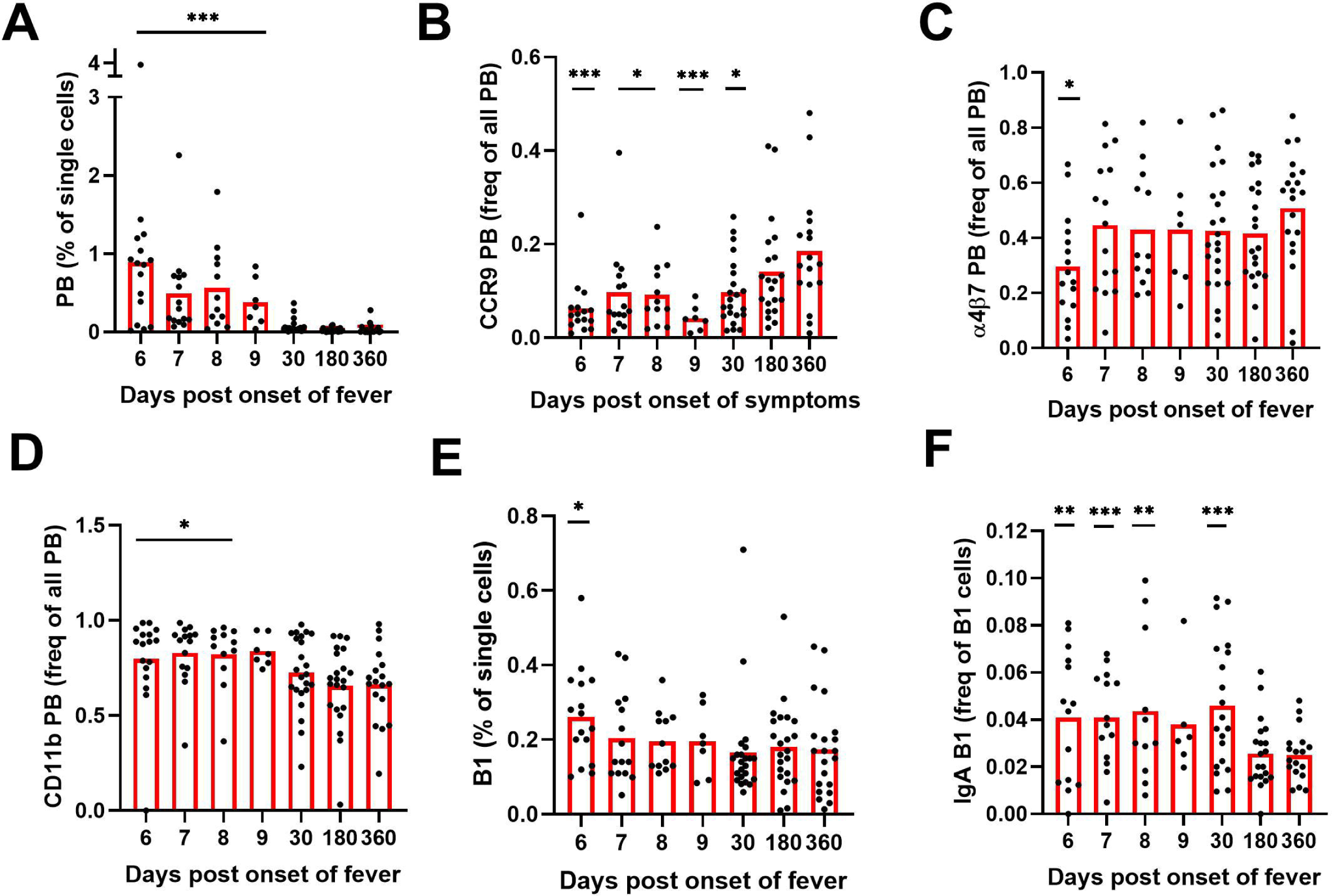
Multiparameter flow cytometric analysis of PBs and B1 cells in peripheral blood of PUUV-HFRS. PBMCs from hospitalized (days 6-9 after onset of fever), two weeks after discharge (∼30 days post onset fever) and recovered (180 and 360 days post-onset of fever) patients (n = 25) were stained with a panel of fluorochrome-labeled antibodies and analyzed by flow cytometry. (A) The frequencies of plasmablasts (PBs) in all single cells. (B) The frequencies of PBs expressing cell surface CCR9. (C) The frequencies of PBs expressing cell surface integrin α4β7. (D) The frequencies of PBs expressing cell surface integrin CD11b. (E) The frequencies of B1 cells in all single cells. (F) The frequencies of surface IgA expressing B1 cells out of all B1 cells. Statistical significance at each time point as compared to recovery at 360 days post-onset of fever were assessed by generalized estimating equations. ***, ** and * indicate p-values <0.001, < 0.01 and < 0.05, respectively.

The frequencies of mucosal-associated B1 cells were significantly increased at 6 days aof as compared to R360 (Fig. 1E). Strikingly however, significantly more B1 cells expressed surface IgA in acute and postacute stages as compared to recovery (approximately 5 % of B1 cells expressed surface IgA at acute stage, Fig 1F). The mucosal homing receptor expressions of α4β7, CD11b, and CCR9 remained unaltered in B1 cells during acute stage as compared to recovery (supplementary Fig. 2E-F), although a slight increase in CCR9 expression was observed at postacute stage. For IgA B1 the cell frequencies were too low for reliable comparisons.

#### IgA ASCs produce mucosal-like IgA during acute PUUV-HFRS

Mucosal IgA is typically polyreactive. To assess the polyreactivity of the IgA produced by the ASCs of the peripheral blood of PUUV-HFRS patients, we cultured PBMCs obtained from patients in acute (1^st^ day of hospitalization) or recovery stage (R360) of the disease for 6 days and analyzed the supernatants for IgA production by ELISA. We utilized a previously established method to assess antibody polyreactivity by measuring its ability to bind dinitrophenol (DNP)-conjugated albumin (Gunti and Notkins 2015) using salivary sIgA as the reference. Consistent with the significantly increased levels of IgA ASCs in acute PUUV-HFRS, we observed markedly higher total IgA production from PBMCs at acute HFRS vs. R360 (Fig. 2A). Interestingly, a notable fraction (∼10-20% estimated using sIgA as a reference) of the IgA produced by PBMCs from the acute stage bound DNP (Fig. 2B), suggesting substantial polyreactivity in the IgA response during acute PUUV-HFRS.

**Figure 2.**
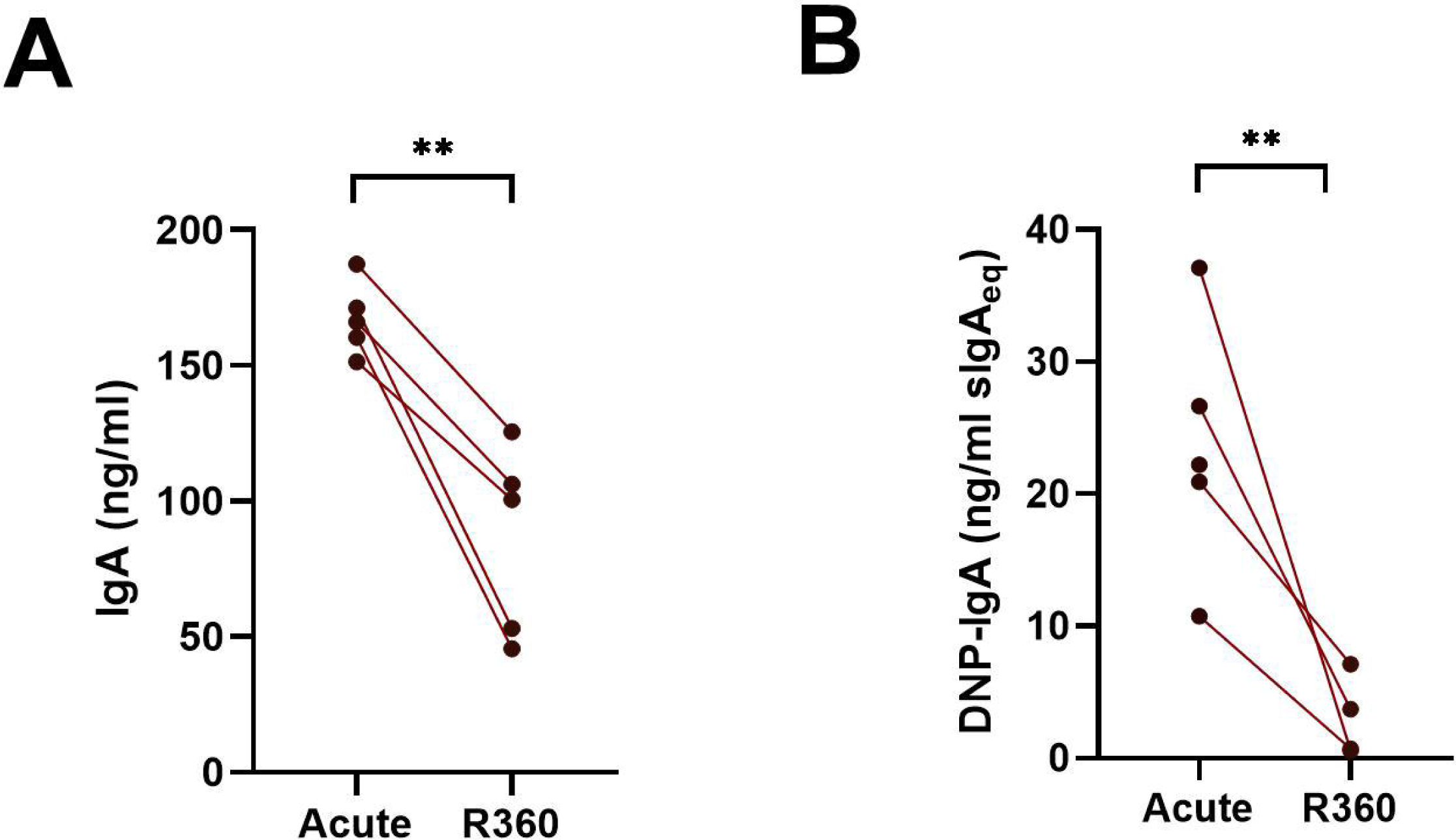
Cells in the peripheral blood spontaneously produce polyreactive IgA during acute PUUV-HFRS. PBMCs isolated from acute (1st day of hospitalization) or recovery (360 days post-onset of fever; R360) of PUUV-HFRS were cultured for 6 days and supernatants analyzed for total IgA (A) and DNP-reactive IgA (B) by ELISA. Salivary sIgA was used as standard in (B) and concentrations reported as sIgA equivalent. The lines connect matched data obtained from the same patient. Statistically significant differences between groups were assessed by paired samples T-test. ** and * respectively indicate p-values <0.01 and <0.05.

#### Increased levels of circulating polyreactive IgA, virus-specific IgA and sIgA in acute PUUV-HFRS

To understand the changes in the composition of circulating IgA during acute PUUV-HFRS, we analyzed sequential serum samples collected during acute PUUV-HFRS (days 5-9 aof), at postacute stage two weeks after discharge (∼30 d aof), and at recovery 180 d and 360 d aof. The total IgA levels did not show significant variation across the time points (Fig. 3A), whereas DNP-reactive IgA was significantly increased during acute illness (5-9 d aof) and remained elevated two weeks after discharge (∼30 d aof) as compared to full recovery at R360 (Fig. 3B). As expected, samples collected during acute illness (5-9 d aof) and in the postacute stage (30 d aof) showed significantly higher virus-specific IgA levels (using PUUV N protein as the antigen) as compared to samples collected following full recovery at R180 and R360 (Fig. 3C).

**Figure 3.**
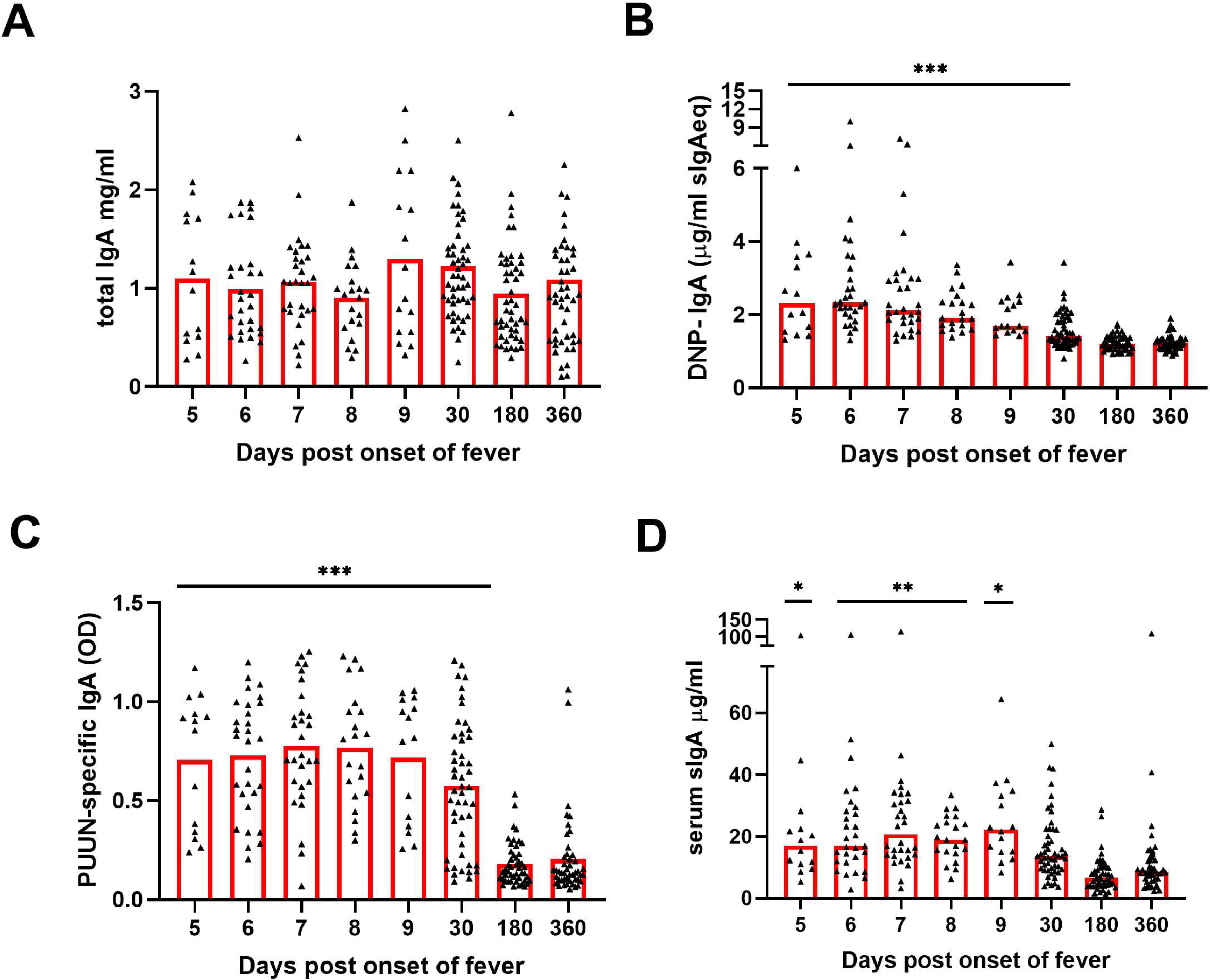
Increased levels of circulating polyreactive, PUUN-specific and secretory IgA in acute PUUV-HFRS. Sequential serum samples from hospitalized (days 5-9 post onset of fever), two weeks after discharge (∼30 days post onset fever) and recovered (180 and 360 days post-onset of fever) PUUV-HFRS patients (n = 55) were analyzed for total IgA (A), DNP-reactive IgA (B), PUUN-specific IgA (C) as well as for secretory IgA (D) by ELISA. Statistically significant differences at each time point as compared to recovery at R360 were assessed by generalized estimating equations. OD = optical density. ***, ** and * indicate p-values <0.001, <0.01 and <0.05, respectively.

To further assess the mucosal origin of the circulating IgA, we developed an ELISA to detect IgA complexed with secretory component (SC), indicative of sIgA. We found significantly increased levels of sIgA in the samples collected in acute phase as compared to those collected in recovery phase of the disease (Fig. 3D). Since SC attaches to IgA only during its transepithelial transport to the mucosal lumen, our findings could be indicative of a breach in the mucosal barrier and leakage of luminal contents into the circulation during acute PUUV-HFRS.

#### IgA autoantibodies to endothelial cells (AECAs) are increased during acute PUUV-HFRS

Endothelial cells (ECs) are the primary target of hantavirus infection, but direct infection of ECs does not typically compromise the monolayer integrity. Therefore, it appears likely that immunological mechanisms contribute to EC permeability and disease progression during HFRS (Mackow and Gavrilovskaya 2009, Hepojoki, Vaheri et al. 2014). We aimed to assess whether circulating IgA could contribute to vascular pathology by binding to infected and/or non-infected ECs during PUUV-HFRS. Patient serum obtained at the first day of hospitalization (acute) or at full recovery (R360) was incubated with PUUV-infected (nearly 100% infection, as shown in Fig. 4A) vs. non-infected (mock) blood microvascular endothelial cells (BECs). After washing, the levels of cell-bound IgA were analyzed by on-cell ELISA. We observed significantly increased IgA binding to both PUUV-infected and mock-infected BECs in acute PUUV-HFRS as compared to R360 (Fig. 4B). Consistent with the presence of virus-specific IgA, the serum obtained from the acute phase showed enhanced binding to infected as compared to mock-infected BECs. The binding of acute-phase IgA to non-infected BECs (in the absence of viral antigen) may be attributed to the previously observed polyreactive nature of IgA, suggesting a virus-independent IgA-mediated immune response towards ECs. This phenomenon could potentially be a contributing factor to the increased EC permeability observed during HFRS.

**Figure 4.**
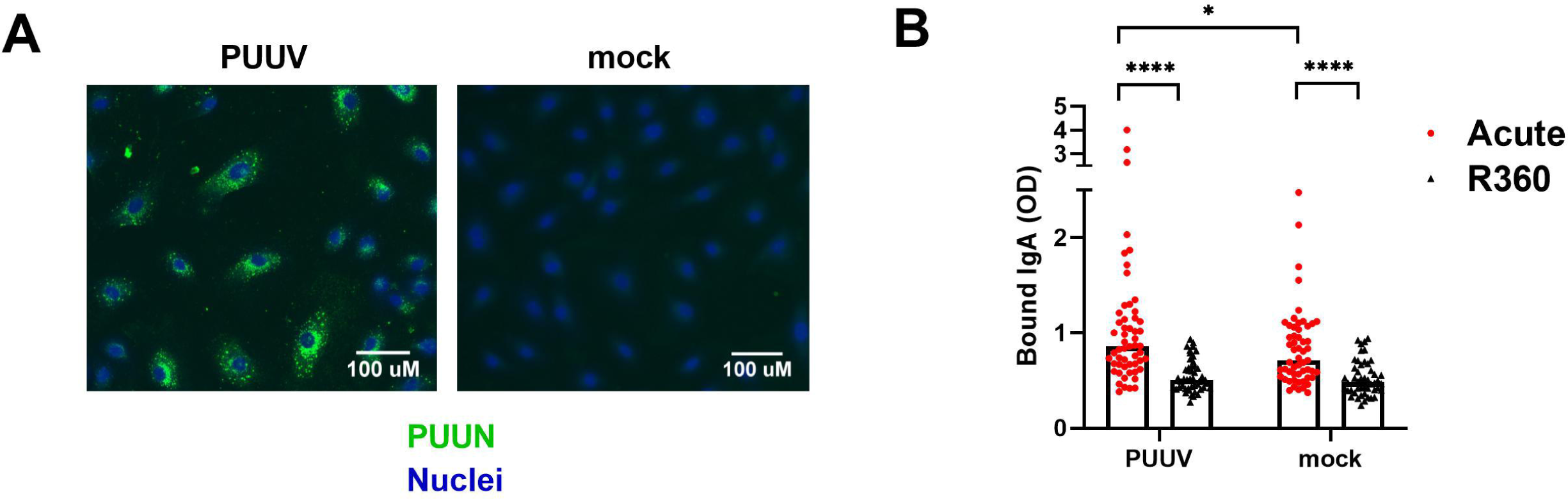
Circulating IgA during acute PUUV-HFRS binds microvascular endothelial cells. PUUV- or mock-infected primary blood microvascular endothelial cells (BECs) were fixed at 24 h post-inoculation and used for studying the IgA responses of acute- and recovery-phase sera. (A) An overlay of immunofluorescence for PUUN (green)- and Hoechst33342 (blue)-stained PUUV- and mock-infected BECs. (B) Serum from acute (1st day of hospitalization) or recovery (360 d post onset of fever) PUUV-HFRS was added on PUUV- or mock-infected BECs for 1 h. After washing, cell-surface bound IgA was measured with on-cell ELISA and optical density (OD) recorded. Statistically significant differences between acute and recovery samples were assessed by two-way ANOVA including Tukey’s multiple comparison test. **** and * indicate p-values <0.0001 and <0.05, respectively.

#### Increased circulating sIgA and virus-specific IgA associate with disease severity

In addition to vascular permeability, the hallmarks of PUUV-HFRS include AKI, the extent of which can be indirectly assessed by measuring blood creatinine levels (Lewington and Kanagasundaram 2011). We stratified the patients based the maximum blood creatinine levels measured during hospitalization as having a mild (blood creatinine ≤ 265 µmol/L, AKI stage 2 or lower) or severe (> 265 µmol/L; AKI stage 3) PUUV-HFRS. Our analyses revealed that the patients with severe illness had higher levels of sIgA and PUUN-specific IgA (Fig. 5A-B), whereas the total IgA, DNP IgA, IgA AECA or IgA ASC levels (Fig. 5C-D and supplementary Fig. 3A-D) did not demonstrate statistically significant differences between the groups. These findings suggest that mucosal and virus-specific IgA responses might play a role in the development of AKI, while IgA polyreactivity *per se* would not. Due to its potential role in vascular pathology, we also stratified patients based on the severity of thrombocytopenia (TC, severe TC with platelet count nadir <50 *10^4^/µl blood) and found that lower platelet counts associated with higher frequency of IgA B1s (Fig. 5E). The levels of IgA, sIgA, N-specific IgA, DNP-IgA, IgA AECA or PB counts appeared to be unaltered between the groups (Fig. 5F-H and supplementary Fig. 3E-H).

**Figure 5.**
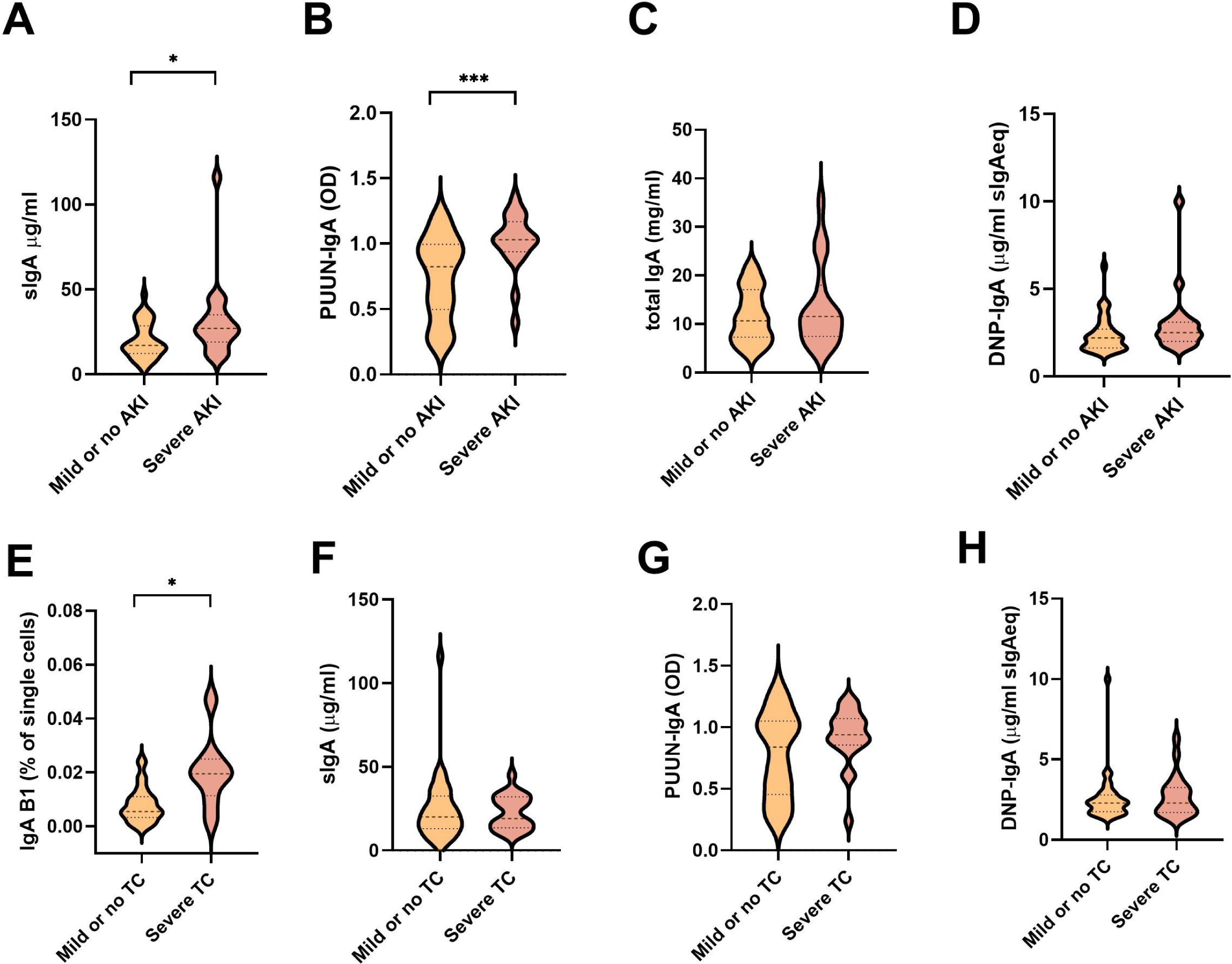
Association between measured soluble IgA and IgA ASC levels with parameters of disease severity. (A-D) The PUUV-HFRS patients were stratified based the maximum blood creatinine levels measured during hospitalization as mild (blood creatinine ≤ 265 µmol/l = AKI stage 2 or lower, n = 36) or severe (blood creatinine > 265 µmol/l = AKI stage 3, n = 19) or (E-H) minimum thrombocyte levels as mild or no thrombocytopenia (TC, ≥ 50 * 10^4^/µl blood) and severe TC (< 50 * 10^4^/µl blood). The maximum levels of sIgA (A, F), PUUN-specific IgA (B, G) DNP-reactive IgA (C, H), total IgA (D), and IgA B1 cells (E) measured during hospitalization were grouped based on patient severity criteria and significant differences assessed by Mann-Whitney test. *** and * respectively indicate p-values <0.001 and <0.05.

#### The polyreactivity and virus neutralization ability of PUUV-induced IgA associates with its multimericity

Secretory IgA appears mostly as dimers (dIgA) but also as higher order complexes (polymeric IgA, pIgA). To understand further the nature of the IgA response and its molecular complexity, we isolated total IgA from acute (1^st^ day of hospitalization) vs. recovery (R360) PUUV-HFRS (patient matched samples, n = 7) and separated the monomeric (mIgA), dimeric (dIgA) and polymeric (pIgA) IgA forms by size-exclusion chromatography (a representative chromatogram and western blot of IgA fractions are shown in supplementary Figure 4A-B). Interestingly, when normalized to mIgA we observed increased levels of dIgA and pIgA in acute vs. R360 samples (Fig. 6A). The mIgA levels were comparable between groups (supplementary Fig. 4C). Thus, multimeric IgA (dIgA and pIgA combined) was approximately 40 % of all circulating IgA in acute PUUV-HFRS when compared to 20% at steady state (R360).

**Figure 6.**
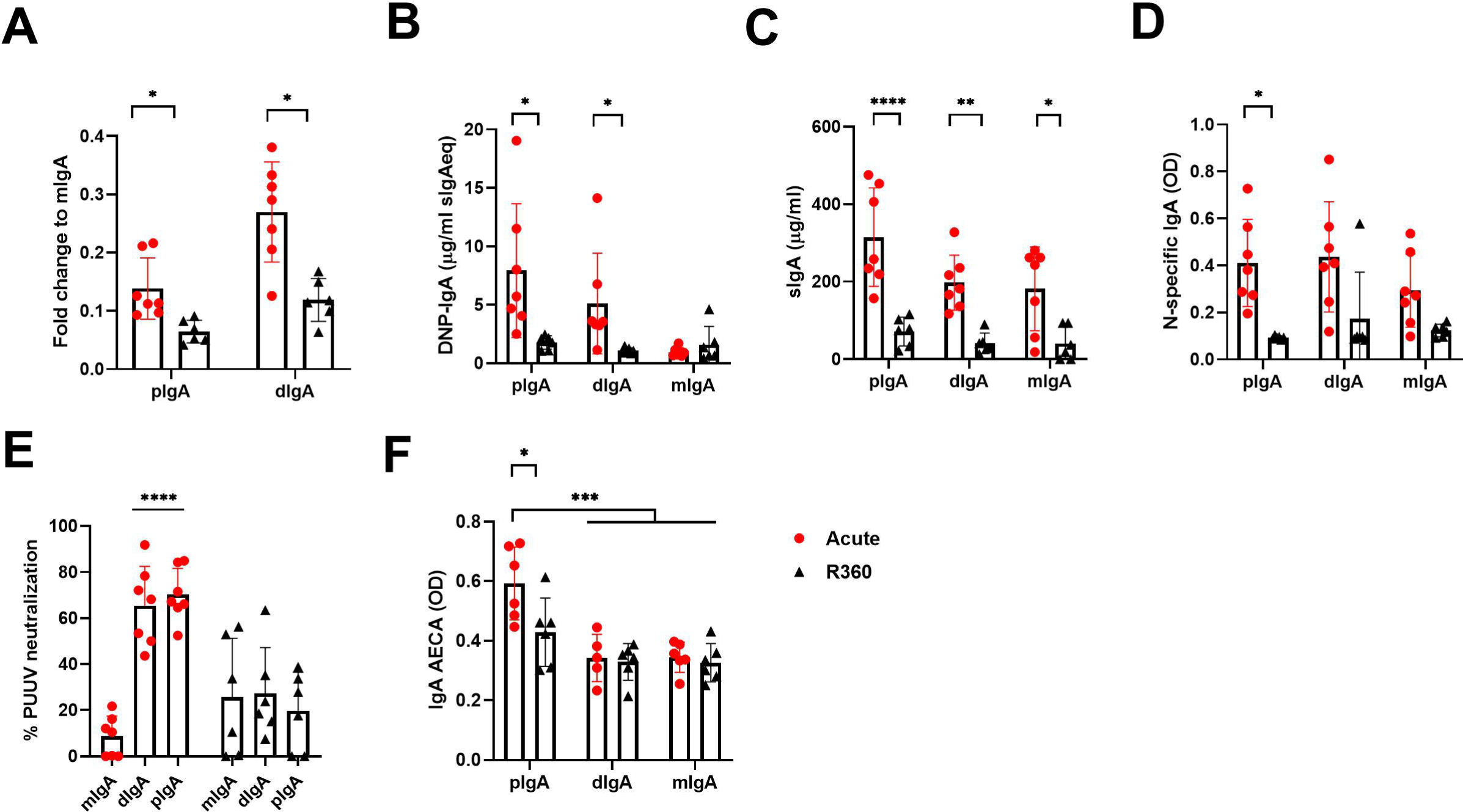
Increased levels of circulating multimeric IgA in acute PUUV-HFRS. Total IgA was isolated from serum of acute (1st day of hospitalization) or recovery (360 d post onset of fever) PUUV-HFRS by Peptide M followed by size –exclusion chromatography to separate polymeric, dimeric and monomeric forms of IgA. Isolated IgA fractions were analyzed for fold change of total IgA concentration compared to mIgA (A). After normalization, IgA fractions were analyzed for DNP-reactive IgA (B), sIgA (C), PUUN-specific IgA (D) and IgA AECAs (F). The neutralization of PUUV was analyzed by a microneutralization test (E). Statistically significant differences between acute and recovery samples of each fraction were assessed by two-way ANOVA including Tukey’s multiple comparison test. ** and * respectively indicate p-values <0.01 and <0.05.

Interestingly, the pIgA and dIgA fractions showed significantly increased polyreactivity and elevated levels of sIgA as compared to recovery phase samples or their mIgA counterparts isolated from the same samples (Fig. 6B-C). However, unexpectedly, increased levels of sIgA were found also mIgA fraction of acute disease (Fig. 6C), possibly indicating breakdown of IgA complexes, while antigen specificity was evenly spread between IgA fractions from acute stage (Fig. 6D). In striking contrast to the latter, PUUV microneutralization assays indicated that pIgA and dIgA, but not mIgA, neutralized virus in cell culture (Fig. 6E). This suggests that efficient virus neutralization requires multimeric IgA. Interestingly, acute pIgA fractions showed elevated EC binding ability as compared to acute dIgA or mIgA, indicating that PUUV-induced IgA AECAs are polymeric (Fig. 6F).

#### Multimeric IgAs form stable complexes with CD89

IgA exerts its effector functions mainly through the IgA receptor CD89. To understand whether the multimericity of IgA in PUUV-HFRS could affect its effector functions, we analyzed the binding kinetics of different IgA fractions to CD89. The isolated IgAs from different donors, as described above, were pooled together in equal ratios and used as analytes in a surface plasmon resonance (SPR) assay, with a CD89-coated surface as the ligand. The kinetic constants were obtained by fitting obtained sensorgrams to a Langmuir model assuming 1:1 binding stoichiometry. All IgA fractions were able to bind to CD89, with mIgA showing strongest apparent dissociation constant Kd_app_ compared to dIgA or pIgA (Table 4 and Figure 7A-C). However, when comparing the affinity rate constants, markedly differing binding kinetics between different IgA fractions were observed. Polymeric IgA exhibited significantly reduced dissociation rates compared to dIgA and mIgA, with mIgA having the highest dissociation rate. This indicated that the complexes formed between dIgA or pIgA and CD89 are more stable compared to those formed by mIgA. On the other hand, mIgA showed significantly increased association rates compared to multimeric IgAs, which translated to its increased overall affinity towards CD89. When comparing IgAs from acute vs. R360 stages of PUUV-HFRS, no significant differences were observed for any IgA fraction (Table 4 and Supplementary Fig. 5A-C).

**Figure 7.**
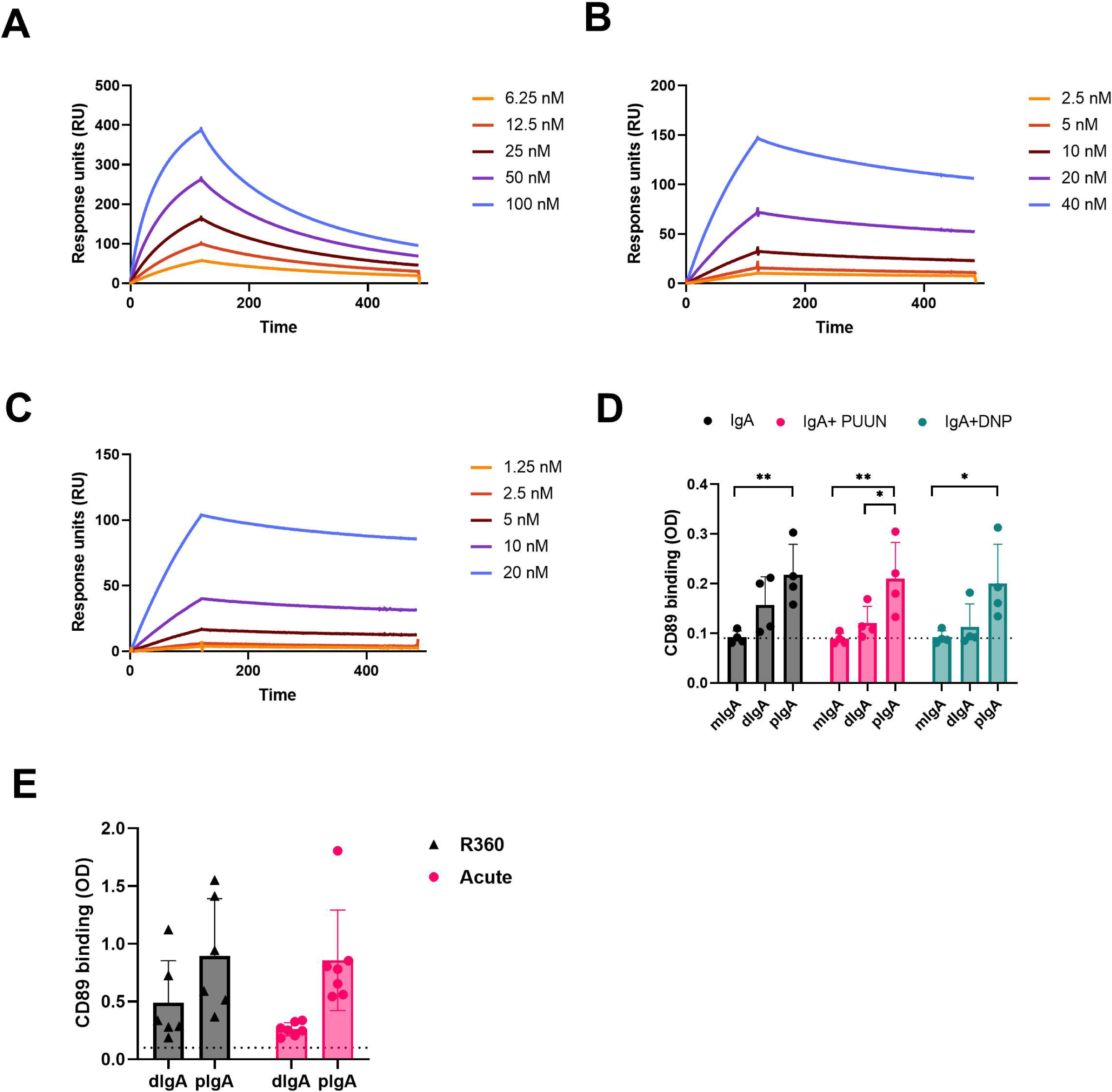
CD89 receptor binding kinetics of circulating mIgA, dIgA and pIgA. (A-C) Surface plasmon resonance (SPR) binding kinetics assay using Biacore T100. The IgA receptor CD89 was coated on the sensor chip surface, and different IgA fractions isolated from acute PUUV-HFRS at indicated concentrations: (A) monomeric IgA (mIgA), (B) dimeric IgA (dIgA) and (C) polymeric IgA (pIgA). (D) CD89 was coated on an ELISA plate, and binding of mIgA, dIgA and pIgA from acute PUUV-HFRS was analyzed. The binding assays were performed in the presence of 100-fold excess PUUN or DNP-albumin where indicated. (E) Comparative CD89-binding ELISA was performed with dIgA and pIgA isolated from acute vs. convalescent PUUV-HFRS. Dotted lines indicate baseline optical density (OD) levels. Statistically significant differences between acute and recovery samples of each fraction were assessed by two-way ANOVA including Tukey’s multiple comparison test. ***, ** and * indicate p-values <0.001, < 0.01 and < 0.05, respectively.

**Table 4.**
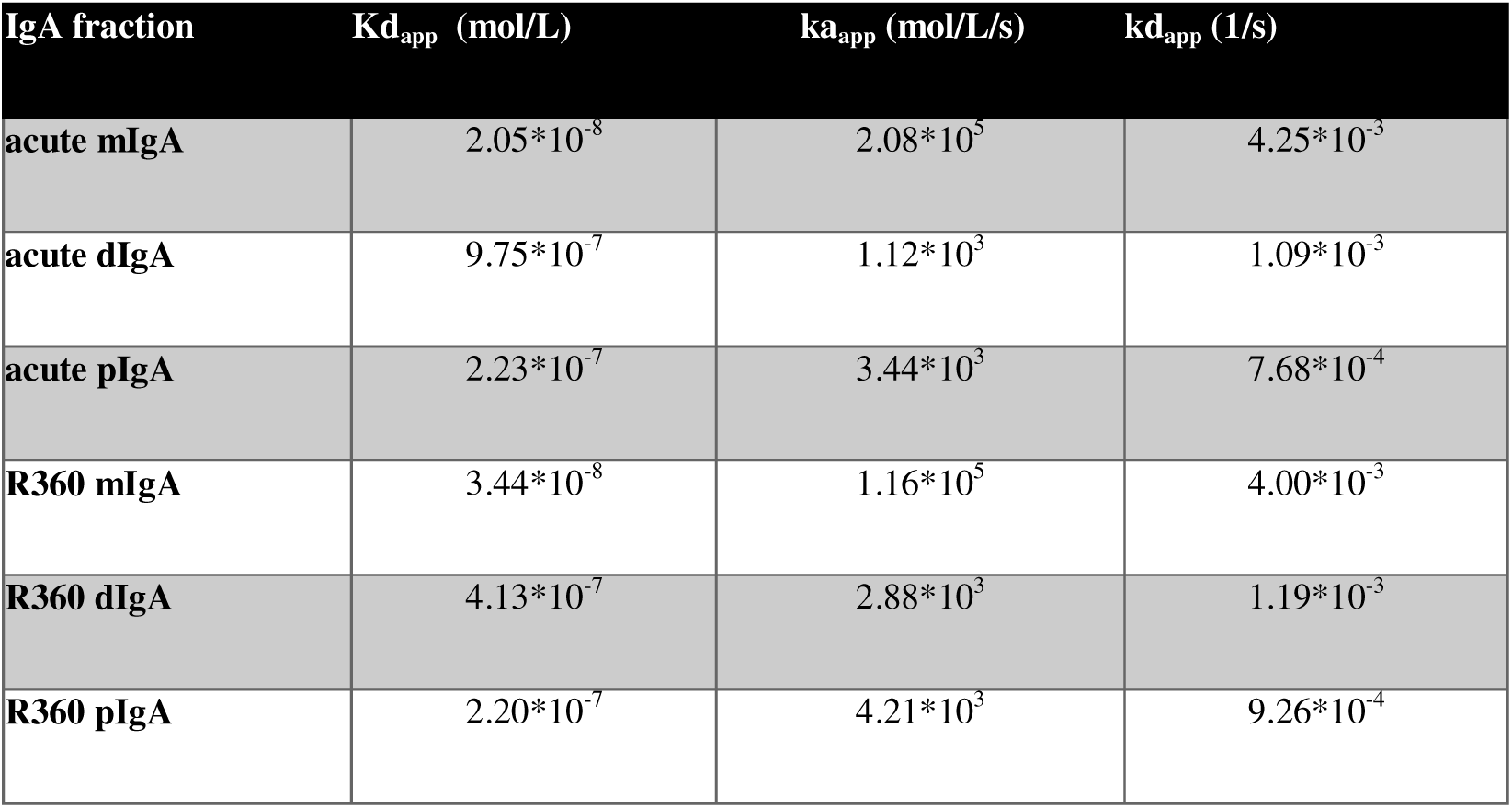
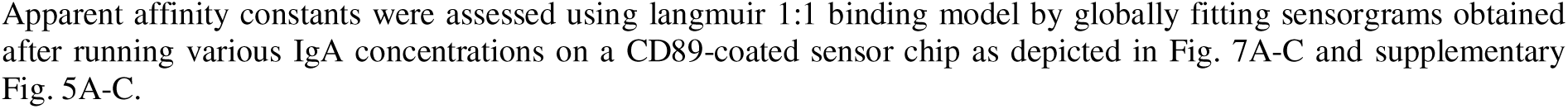
Apparent kinetic affinity constants between IgA fractions and the IgA receptor CD89.

| IgA fraction | K<sub>d,app</sub> (mol/L) | k<sub>a,app</sub> (mol/L/s) | k<sub>d,app</sub> (1/s) |
| --- | --- | --- | --- |
| **acute mIgA** | 2.05*10<sup>-8</sup> | 2.08*10<sup>5</sup> | 4.25*10<sup>-3</sup> |
| **acute dIgA** | 9.75*10<sup>-7</sup> | 1.12*10<sup>3</sup> | 1.09*10<sup>-3</sup> |
| **acute pIgA** | 2.23*10<sup>-7</sup> | 3.44*10<sup>3</sup> | 7.68*10<sup>-4</sup> |
| **R360 mIgA** | 3.44*10<sup>-8</sup> | 1.16*10<sup>5</sup> | 4.00*10<sup>-3</sup> |
| **R360 dIgA** | 4.13*10<sup>-7</sup> | 2.88*10<sup>3</sup> | 1.19*10<sup>-3</sup> |
| **R360 pIgA** | 2.20*10<sup>-7</sup> | 4.21*10<sup>3</sup> | 9.26*10<sup>-4</sup> |

To confirm the observed differences using pooled IgA fractions, we set up an ELISA assay to analyze the binding of individual patient IgAs to CD89-coated plates. This assay allowed us to measure IgA-CD89 interactions in the presence of the previously identified antigens PUUN and DNP-albumin. Interestingly, compared to dIgA or mIgA, pIgA showed significantly increased binding to CD89, while the presence of excess amounts of antigen did not significantly alter the interaction efficiency in this assay (Fig. 7G). These findings suggest that the increased stability of pIgA-CD89 interaction, as observed by the kinetic analysis using SPR, plays a major role in the outcome of this ELISA assay. However, the presence of antigen did not impact IgA binding to CD89, indicating that the interaction is mediated by the Fc portion of IgA, as expected. Similarly to the SPR analysis, no significant differences were observed between IgAs isolated from the acute vs. R360 phases in the ELISA (Fig. 7E).

#### pIgA activates neutrophils

The IgA receptor CD89 is widely expressed in neutrophils, which suggests that neutrophils could play a key role in sensing changes in the IgA composition during PUUV-HFRS. We therefore analyzed whether the isolated IgA fractions could mediate neutrophil responses. We isolated neutrophils from healthy volunteers and incubated the isolated neutrophils with different IgA fractions for 4-hr before being analyzed for reactive oxygen species (ROS) generation and expression of surface markers LOX-1, CD66b, CD11b, CD62L and HLA-DR by flow cytometry.

Indeed, irrespective of the sampling time point (acute or R360), the pIgA differed significantly from dIgA or mIgA in its ability to induce increased frequencies of ROS producing as well as LOX-1 and high CD66b expressing neutrophils (Fig. 8A-C, representative histograms shown in supplementary Figure 6). Furthermore, the pIgA-treated neutrophils displayed lower frequency of CD62L expression (Fig. 8D) and increased expression of CD11b and HLA-DR as judged by elevated MFI (Fig. 8E-F). Collectively, these data indicate that pIgA activates pro-inflammatory responses in neutrophils, which could be linked to its ability to form stable complexes with CD89. Importantly, the pro-inflammatory effect of pIgA did not significantly differ between acute vs. R360 of PUUV-HFRS, suggesting that it is an intrinsic characteristic of pIgA and not directly related to acute disease. However, due to the significantly increased levels of pIgA in acute PUUV-HFRS it is likely that net pro-inflammatory effect of pIgA is significantly elevated as compared to steady state.

**Figure 8.**
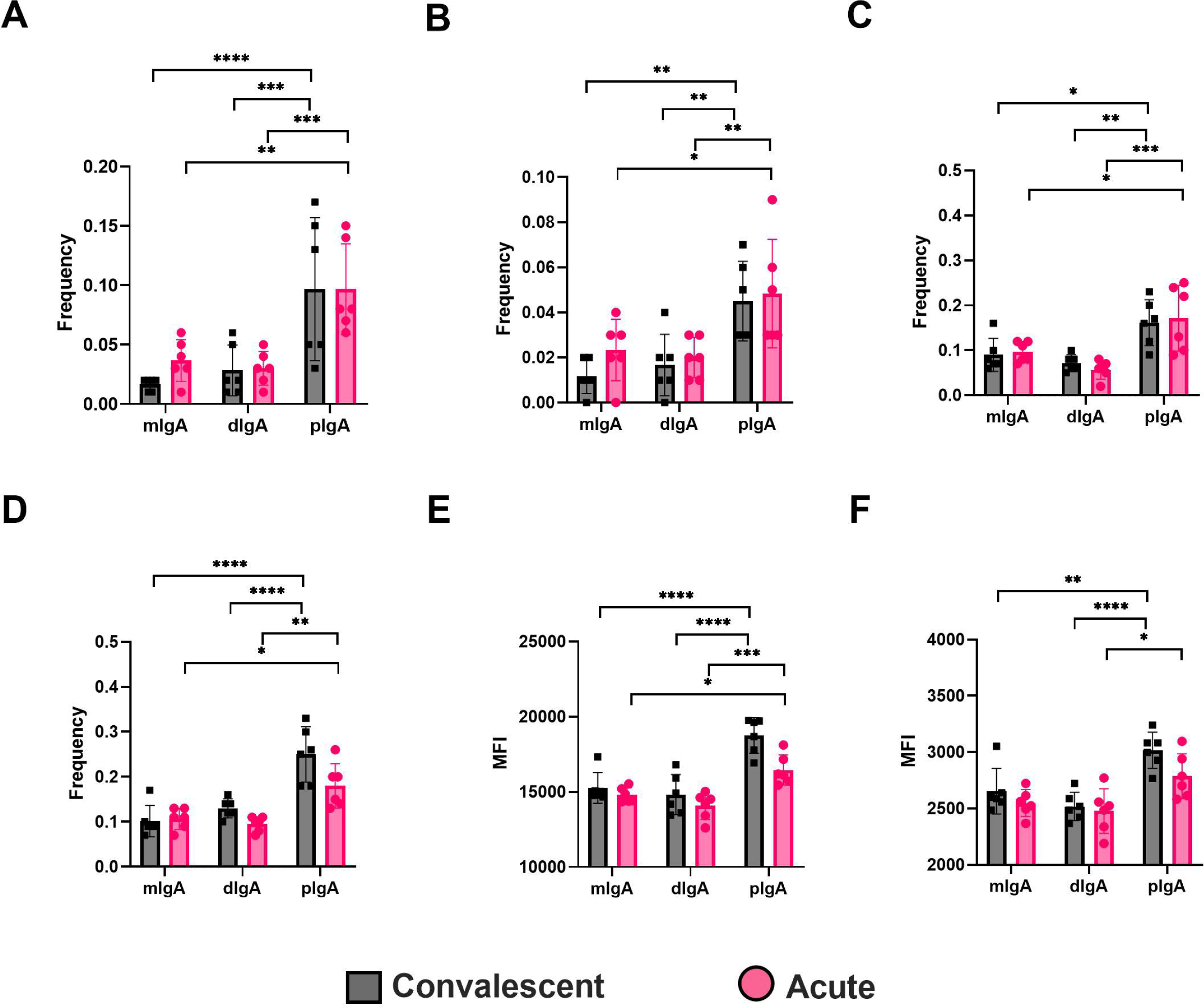
Circulating pIgA activates neutrophils. Neutrophils were isolated from healthy volunteers and incubated 4 h with 10 µg/ml of pIgA, dIgA and mIgA obtained from acute vs. recovery PUUV-HFRS patients as previously described. Reactive oxygen species (ROS) generation (A) and cell surface expressions of LOX-1 (B), CD66b (C), CD62L (D), CD11b (E) and HLA-DR (F) were assessed by multicolor flow cytometry. The frequency of gated positive cells (A-C), negative cells (D) or median fluorescence intensity (MFI) of all cells (E, F) were reported. Statistically significant differences between acute and recovery samples of each fraction were assessed by two-way ANOVA including Tukey’s multiple comparison test. ****, ***, ** and * respectively indicate p-values <0.0001, <0.001, <0.01 and <0.05, respectively.

## Discussion

The pathogenesis of HFRS is mediated by an exacerbated immune response to the causative virus (Klingström, Smed-Sörensen et al. 2019). The early phase of orthohantavirus infections in humans remains unclear, but since it is acquired through inhalation from rodent excreta, the virus must initially make contact with host mucosal compartments. However, the role of mucosal immunity in hantavirus-mediated diseases remains obscure. Mucosal immunity is largely mediated by IgA, which, given the large mucosal surface area, is the most abundant immunoglobulin in humans. Previous reports indicate that circulating IgA responses are elevated in acute PUUV-HFRS (de Carvalho Nicacio, Björling et al. 2000) and that a majority of PBs induced by orthohantavirus infection express IgA in both in HCPS and HFRS (García, Iglesias et al. 2017, Hepojoki, Cabrera et al. 2021). These findings imply that the mucosal immune compartment is activated by orthohantavirus infection, but the exact type of induced IgA and its role in disease progression are not well understood.

In this study, we analyzed the IgA-expressing cellular landscape in the periphery in more detail. In addition to the IgA PBs previously described, we found that circulating IgA-expressing B1 cells are also significantly increased and likely contribute to the total IgA response in acute PUUV-HFRS. Consistent with the upregulation of peripheral IgA ASCs, we detected increased levels of polyreactive and secretory IgA in blood of patients. Notably, the elevated levels of sIgA were associated with worsened AKI, a hallmark of HFRS. Furthermore, we found that infection-induced IgA interacts with endothelial cells and circulates as multimeric complexes, which strongly associated with the IgA receptor CD89 and induced pro-inflammatory responses in neutrophils. These results suggest a model in which PUUV infection induces aberrant translocation of pro-inflammatory mucosal IgA responses to the periphery, potentially contributing to the disease progression of PUUV-HFRS.

Plasma cell trafficking is mediated by adhesion molecules and chemokine receptors (Kunkel and Butcher 2003). The integrins α4β7 and α4β1 respectively mediate the migration of plasma cells and PBs to the intestine and non-intestinal sites, such as the respiratory tract. Additionally, intestinal IgA plasma cells are characterized by CD11b expression (Kunisawa, Gohda et al. 2013) while CCR9 directs cells specifically to the small intestine (Hieshima, Kawasaki et al. 2004). In the current study, we investigated the surface expression of α4β7, CD11b and CCR9 in infection-induced PBs to better understand their migratory patterns in mucosal versus systemic compartments. Since plasma cells in the steady state express high levels of the intestinal homing receptor α4β7 (Mei, Yoshida et al. 2009) they serve as a reference phenotype for intestinal PBs. We found that infection-induced PBs displayed significantly less CCR9 expression throughout the acute phase (hospitalization), while α4β7 was also downregulated at the early time point compared to PBs in the patient-matched recovered steady state. In contrast, CD11b expression was strongly elevated in acute PBs. Thus, our findings suggest that infection-induced PBs are distinct from the typical mucosal PBs and, based on their low CCR9 expression, are not targeted to the small intestine. However, with the exception of the early time point at 6 d aof, the expression of α4β7 in PBs during acute disease was comparable to that in the steady state, suggesting that a significant proportion of acute PBs home to the intestine, while other compartments may also be targeted in the early phases of the disease. For instance, some PBs could target the kidneys, where we previously observed significant plasma cell infiltration in acute PUUV-HFRS (Hepojoki, Cabrera et al. 2021). The increased expression of CD11b also supports the intestinal homing of infection-induced PBs and may reflect their vigorous proliferation and increased IgA expression, as previously described for CD11b+ IgA plasma cells over their CD11b-counterparts (Kunisawa, Gohda et al. 2013, Fu, Wang et al. 2021). In light of these findings, we were surprised to detect diminished frequencies of IgA+PBs in the late acute stage (day 9 aof) compared to the steady state. However, this is probably due to increased IgA secretion by IgA+ PBs, leading to reduced IgA levels expressed on the cell surface.

B cells can be divided into the conventional B cells (B2) and “innate-like” B1 cells. While B2 cells generate specific antibody responses against foreign antigens, typically involving T cell–dependent affinity maturation and somatic hypermutation, B1 cells are predominantly found in the peritoneal and pleural cavities in mice, where they produce natural antibodies, providing a “first line of defense” against infections. B1 cells are relatively poorly studied in humans, with controversy existing regarding the exact phenotype that defines human B1 cells (Suchanek and Clatworthy 2023). Despite this, due to their residency in mucosal compartments and ability to quickly respond to infections, we were interested in whether these cells could also contribute to mucosal IgA responses during PUUV-HFRS. We defined human B1 cells as CD20^+^CD27^+^CD38^-/int^CD43^+^CD70^-^(Quách, Rodríguez-Zhurbenko et al. 2016) and found significantly elevated circulating levels of surface IgA-expressing B1 cells in acute disease, while the total number of B1 cells was also slightly elevated. These findings suggest that, in addition to IgA PBs/plasma cells, B1 cells also contribute to the total IgA ASC responses during PUUV-HFRS. A large fraction of B1 cells have been described to produce natural IgM antibodies in mice (Suchanek and Clatworthy 2023). Our study supports the notion that most circulating human B1 cells express IgM, but their frequencies are not similarly upregulated as compared to IgA B1s. Furthermore, we found that a substantial portion of B1 cells expressed α4β7, CD11b and CCR9. However, no changes in the frequencies of these mucosal homing receptors were observed between acute PUUV-HFRS and steady state for total B1 cells. Unfortunately, the low overall low frequencies of IgA B1 cells did not allow for reliable receptor expression comparisons between acute infection and steady state.

We wanted to determine whether the elevated mucosal-type IgA ASCs could significantly contribute to the total circulating IgA pools. Since mucosal IgA is polyreactive, we made use of a previously established method to assess antibody polyreactivity through its binding to DNP. DNP is a synthetic molecule not present in the environment, and individuals are not normally exposed to it. It has been shown previously for IgG that antibodies that bind DNP can be considered polyreactive, and their titer to DNP could serve as an index of polyreactivity (Gunti and Notkins 2015). Thus, in the current study we applied this approach to measure the polyreactivity of IgA. We observed that *in vitro* cultivated PBMCs from acute PUUV-HFRS spontaneously released significant levels of DNP-reactive IgA, which was strongly elevated along also in the serum of patients with acute disease, alongside the expected virus-specific IgA responses. Further supporting the involvement of mucosal IgA responses in acute PUUV-HFRS, we also observed increased levels of sIgA in circulation. sIgA is typically found only in mucosal secretions, to which it enters through transepithelial transport. This suggests a breach of the mucosal layers and leakage of its luminal contents into the periphery. Thereby, our findings indicate that increase in the circulating mucosal-type IgA in acute PUUV-HFRS would be due to both the increased frequency of IgA ASCs and increased mucosal permeability.

Alterations in the EC permeability are characteristic of the orthohantavirus-mediated diseases, with ECs being the prime target of virus replication. Considering the observed IgA polyreactivity and virus specificity, we assessed whether circulating IgA can bind infected and non-infected ECs. Indeed, we found increased levels of IgA bound to *in vitro* PUUV infected and non-infected ECs compared to IgAs from steady state controls. Since the binding was also observed to non-infected ECs, this suggests that IgA polyreactivity may play role in this phenomenon. Importantly, the presence of IgA AECAs in acute PUUV-HFRS indicates that IgA may play role in the development of HFRS pathogenesis by attracting immune responses towards ECs, which could be a causative factor behind increased EC permeability. Interestingly, mucosal IgA is known for its ability to induce the alternate pathway of complement activation (Hiemstra, Gorter et al. 1987), which is significantly induced in PUUV-HFRS (Sane, Laine et al. 2012). Thereby it is plausible that IgA AECAs could induce complement activation on the surface of both infected and non-infected ECs.

Mucosal Ig is predominantly found in dimers (dIgA) but also forms higher order complexes (pIgA). In line with the observed increase in systemic mucosal IgA responses, we detected increased levels of circulating dIgA and pIgA in acute PUUV-HFRS. IgA complex formation is known to influence its effector functions by facilitating cross-linking of the IgA receptor CD89 (Breedveld and van Egmond 2019). Therefore, we investigated the interaction kinetics between the different IgA forms (mIgA, dIgA and pIgA) and CD89 using surface plasmon resonance technology. Interestingly, both dIgA and pIgA differed significantly from mIgA in their slower affinity rate constants. The differences in the dissociation rates were particularly notable, with pIgA showing the slowest rate followed by dIgA, potentially influencing downstream effector functions by enhancing receptor cross-linking. While previous studies have reported the slower dissociation rate of pIgA over mIgA (Oortwijn, Roos et al. 2007), the slower association rates of dIgA and pIgA observed in this study were unexpected, resulting in an overall stronger binding affinity for mIgA. It is important to mention that although the binding between CD89 and mIgA has been shown to follow 2:1 stoichiometry (Herr, White et al. 2003), we reported the apparent affinity based on a 1:1 binding model for simplicity. Interestingly, our ELISA binding experiments between IgA and CD89 reflected the differences determined by their variable dissociation rates (pIgA > dIgA > mIgA), suggesting that the reduced association rate of multimeric IgAs was not a limiting factor in the ELISA assay setup. Importantly, IgAs isolated from the convalescent stage of the disease (R360) showed similar affinities to CD89 as IgAs from acute stage, implying that the observed differences between IgA fractions were indeed due to their variable degrees of intrinsic complexity and not related to their antigen specificity or polyreactivity.

In addition to the known pro-inflammatory effects of multimeric IgA (Breedveld and van Egmond 2019), we observed elevated ROS generation and increased expression of LOX-1, CD66b, CD11b and HLA-DR in neutrophils upon stimulation with pIgA, but not with dIgA or mIgA. Similarly as with its interaction with CD89, the source of pIgA (whether isolated from acute or steady state) did not significantly influence its pro-inflammatory effect, suggesting that it is independent of antigen specificity and rather an intrinsic property of pIgA. While the increased pro-inflammatory effects of pIgA are associated with its slower dissociation rates from CD89, a receptor strongly expressed on neutrophils (Breedveld and van Egmond 2019), we cannot exclude the possibility that other receptors wmay play an accessory role in IgA-mediated effector functions in neutrophils. For instance, the effector functions of sIgA are mediated by its binding to Mac-1 on neutrophils (Van Spriel, Leusen et al. 2002).

We investigated the potential impact of circulating IgA ASCs and soluble IgAs on the clinical severity of PUUV-HFRS by stratifying patients based on two independent factors reflecting disease severity: acute kidney injury (AKI) and thrombocytopenia. Remarkably, patients with severe AKI (stage 3) showed significantly elevated levels of sIgA and PUUN-specific IgA in circulation. The link between mucosal IgA responses and kidney disease is widely recognized in the case of IgA nephropathy (Seikrit and Pabst 2021). Interestingly, while the pathology of IgA nephropathy has been linked to aberrantly glycosylated IgA, increased sIgA deposits in the kidneys have also been observed (Oortwijn, Rastaldi et al. 2007). Whether similar pathological mechanisms contribute to renal complications in HFRS warrants further investigation. Regarding thrombocytopenia, patients with more severe platelet depletion had a higher frequency of IgA B1 cells. This finding suggests a possible connection between mucosal IgA responses and thrombocytopenia during acute PUUV-HFRS. However, the lack of association between soluble IgA levels and severe thrombocytopenia suggests that this interplay may involve additional factors beyond the release of IgA from B1 cells.

Our findings, demonstrating a robust association between the magnitude of virus-specific IgA response and AKI, align with our previous studies indicating a strong association between circulating antibody light chain levels and kidney dysfunction in PUUV-HFRS (Hepojoki, Cabrera et al. 2021). This supports the concept of immunologically mediated mechanisms contributing to AKI during PUUV-HFRS. Interestingly, a similar strong association between PUUN-specific IgG levels and AKI was not found (Iheozor-Ejiofor, Vapalahti et al. 2022). Our study also revealed a potential mechanism by which IgA could exacerbate disease progression by triggering pro-inflammatory responses in neutrophils, whether there are elicited on the surfaces of infected and non-infected ECs, or possibly on other cells of the kidney. Our data asupports the hypothesis that PUUV infection initiates a strong mucosal IgA response, which then passively “leaks” into the circulation, where its excessive pro-inflammatory effects engage innate immune responses.

Our study has some limitations worth discussing. First, our reliance on blood samples from PUUV-HFRS patients restricted the assessment of mucosal IgA responses solely to systemic circulation. Analyzing the type and specificity of mucosal IgA responses in feces, urine and saliva during acute PUUV-HFRS would provide a more comprehensive understanding of the complete IgA response towards PUUV infection. Secondly, our study did not directly address the interplay between IgA polyreactivity and antigen specificity. Understanding whether one IgA clone can simultaneously exhibit antigen-specific, polyreactivity and covalently linked to SC would be beneficial. Additionally, while our findings are suggestive of a pathological role for IgA in PUUV-HFRS, further studies are needed to elucidate the precise mechanisms through which IgA contributes to the development of AKI, thrombocytopenia and vascular permeability.

## Supplemental figure legends

**Supplementary Figure 1. Gating strategy allowing for the identification of PBs and B1 cells in PBMC of PUUV-HFRS patients by flow cytometry**. After gating of PBMCs (SSC-H vs. FSC-H), single cells (SSC-H vs. SSC-A) and live cells (excluding also non-B cells expressing CD3, CD14, CD56 and CD66), B cells were identified as positive for either CD19, CD20 or both. PBs were identified as CD20±CD19+IgD-CD27+CD38++CD43+ cells and B1 cells as CD20+CD27+CD38-/intCD43+CD70-. The cells positive for surface expression of IgA, integrin α4β7, CD11b or CCR9 in PBs and B1 were gated as indicated. Contour plots are shown.

**Supplementary Figure 2. Multiparameter flow cytometric analysis of IgA PBs and B1 cells in peripheral blood of PUUV-HFRS**. PBMCs from hospitalized (days 6-9 post onset of fever), two weeks after discharge (∼30 days post onset fever) and recovered (180 and 360 days post-onset of fever) patients (n = 25) were stained with a panel fluorochrome-labeled antibodies and analyzed by flow cytometry. (A) The frequencies of PBs expressing cell surface IgA. (B) The frequencies of IgA PBs expressing surface integrin α4β7. (C) The frequencies of IgA PBs expressing surface integrin CD11b. (D) The frequencies of B1 cells expressing surface integrin α4β7. (E) The frequencies of B1 cells expressing surface CD11b. (F) The frequencies of B1 cells expressing surface CCR9. Statistically significant differences at each time point as compared to recovery at 360 days post-onset of fever were assessed by generalized estimating equations. ***, ** and * indicate p-values <0.001, < 0.01 and < 0.05, respectively.

**Supplementary Figure 3. Association between measured soluble IgA and IgA ASC levels with parameters of disease severity**. (A-D) The PUUV-HFRS patients were stratified based the maximum blood creatinine levels measured during hospitalization as mild (blood creatinine ≤ 265 µmol/l = AKI stage 2 or lower, n = 36) or severe (blood creatinine >265 µmol/l = AKI stage 3, n = 19) or (E-H) minimum thrombocyte levels as mild or no thrombocytopenia (TC, ≥ 50 * 10^4^/µl blood) and severe TC (< 50 * 10^4^/µl blood). The maximum levels of PUUV-EC binding IgA (A, F), IgA AECA (B, G), PBs (C, F), IgA B1 cells (D) and total IgA (E) measured during hospitalization were grouped based on patient severity criteria and significant differences assessed by Mann-Whitney test.

**Supplementary Figure 4. Isolation of mIgA, dIgA and pIgA from serum of PUUV-HFRS patients**. (A) Representative chromatogram of the separation of pIgA, dIgA and mIgA from total IgA isolated from patient serum. (B) Representative non-reducing western blot indicating the molecular weight of isolated mIgA (∼180 kDa), dIgA (∼360 kDa) and pIgA (> 360 kDa) fractions. The IgA bands were visualized by a polyclonal anti-kappa light chain antibody followed by IRdye-conjugated secondary antibody. Molecular size was estimated using as molecular weight standard on separate lane. (C). The IgA concentration of isolated IgA fractions from acute and R360 phase of PUUV-HFRS (n = 7 for both) were measured by total IgA ELISA.

**Supplementary Figure 5. CD89 receptor binding kinetics of circulating mIgA, dIgA and pIgA isolated from convalescent PUUV-HFRS analyzed by surface plasmon resonance**. The IgA receptor CD89 was coated on the surface of biacore sensor chip and different IgA fractions, (A) mIgA, (B) dIgA and (C) pIgA, isolated from recovery PUUV-HFRS (360 days post onset of fever) at indicated concentrations were used as analytes in a surface plasmon resonance binding kinetics assay using Biacore T100.

**Supplementary Figure 6. Representative histograms of pro-inflammatory marker expression in neutrophils.** Neutrophils were isolated from healthy volunteers and incubated mIgA, dIgA and pIgA isolated from PUUV-HFRS patient serum at acute vs. recovery stages of the disease. The expression of indicated markers were assessed by flow cytometry and representative histograms of pIgA (solid line with grey area)- vs. mIgA (dotted line with white area)-treated neutrophils are shown. The gating area to select ROS (A) and LOX-1 (B) positive cells as well as cells with high CD66b expression (C) is indicated. The cells with diminished CD62L expression were selected by gating as indicated in (D). The increased expression of CD11b (E) and HLA-DR (F) were assessed by Median fluorescence intensity (MFI).

