## Supplementary figures and images for "Circulating mucosal-like IgA responses associate with severity of Puumala orthohantavirus-caused hemorrhagic fever with renal syndrome"

### Suppl Fig.1

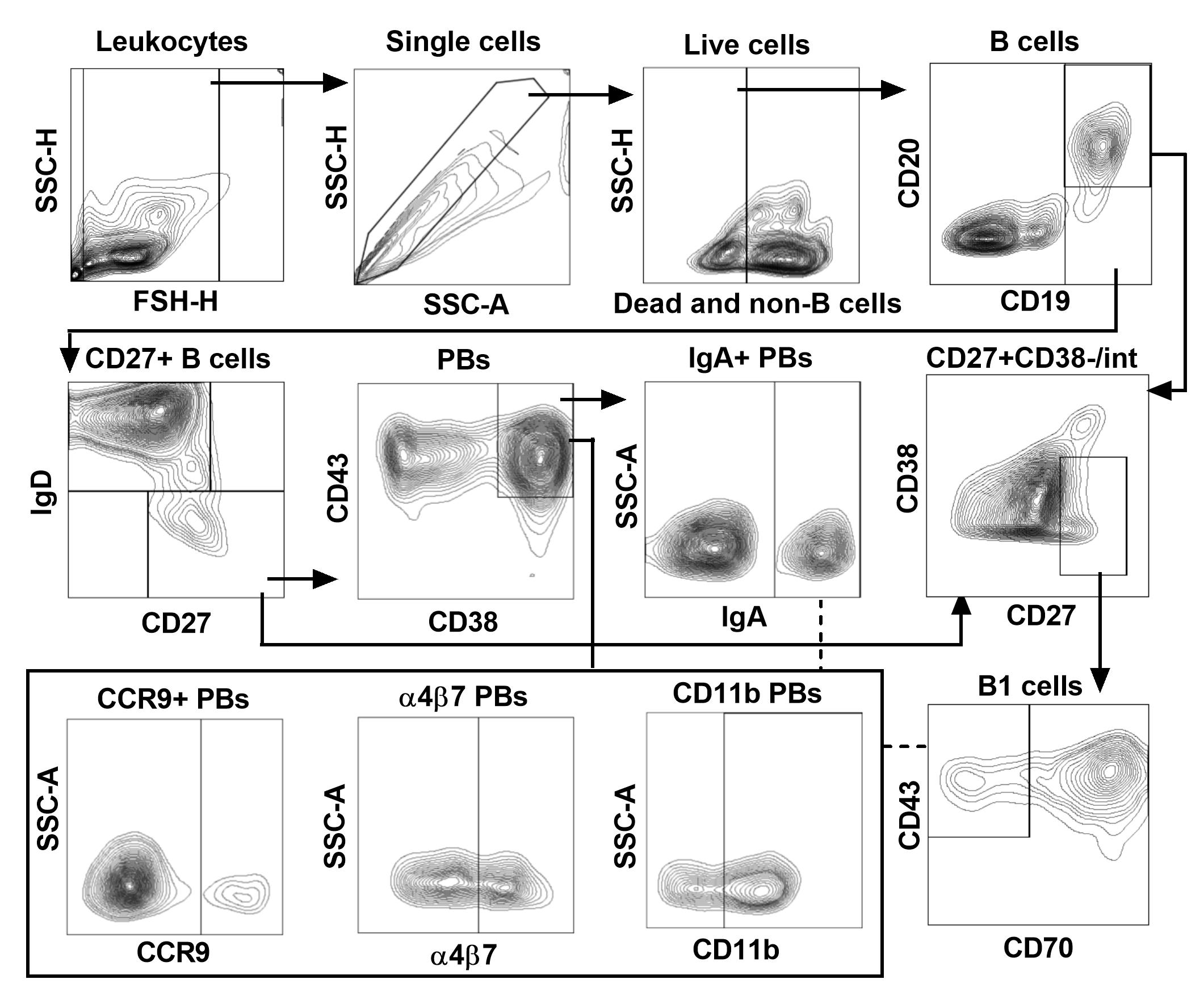

### Suppl Fig.2

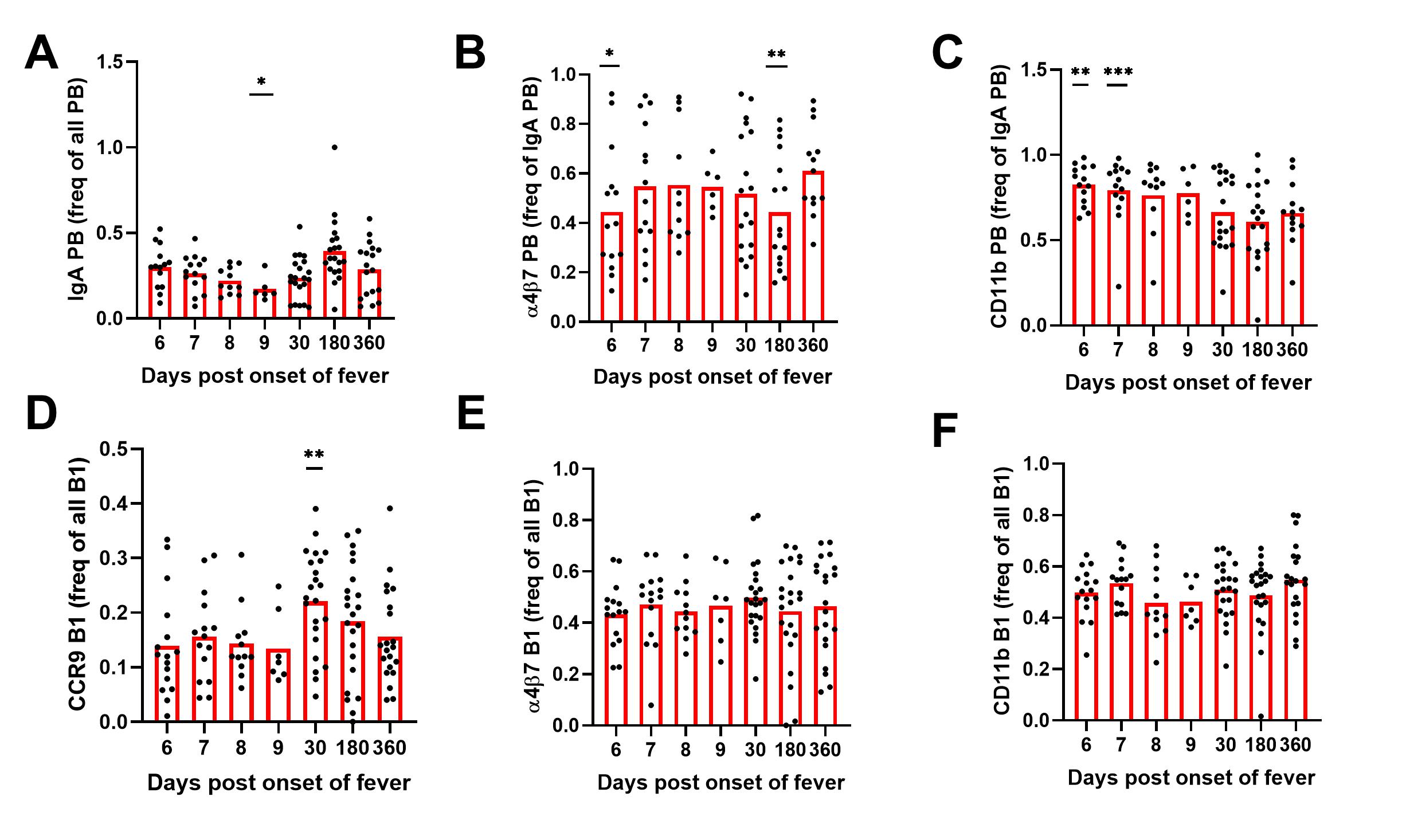

### Suppl Fig.3

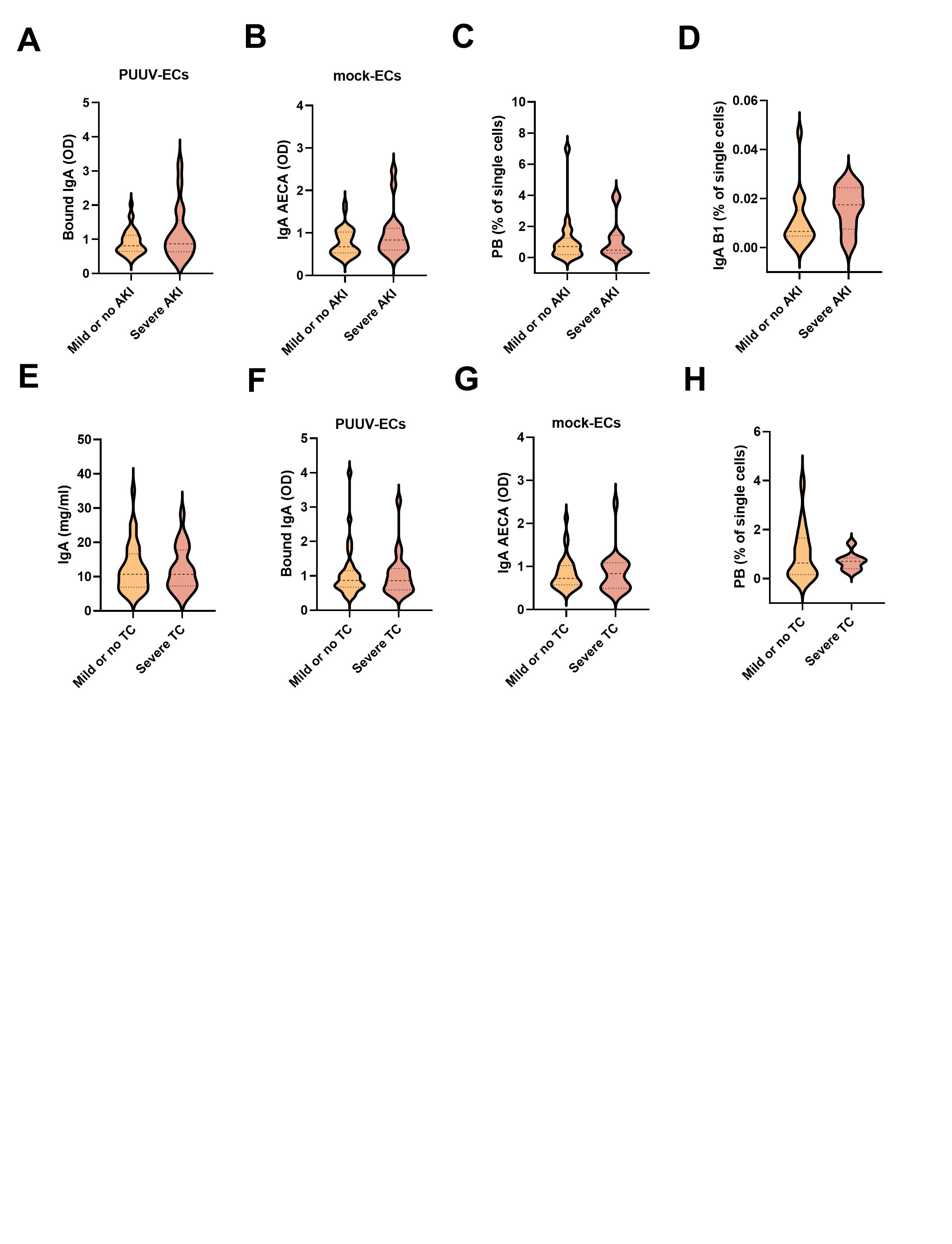

### Suppl Fig.4

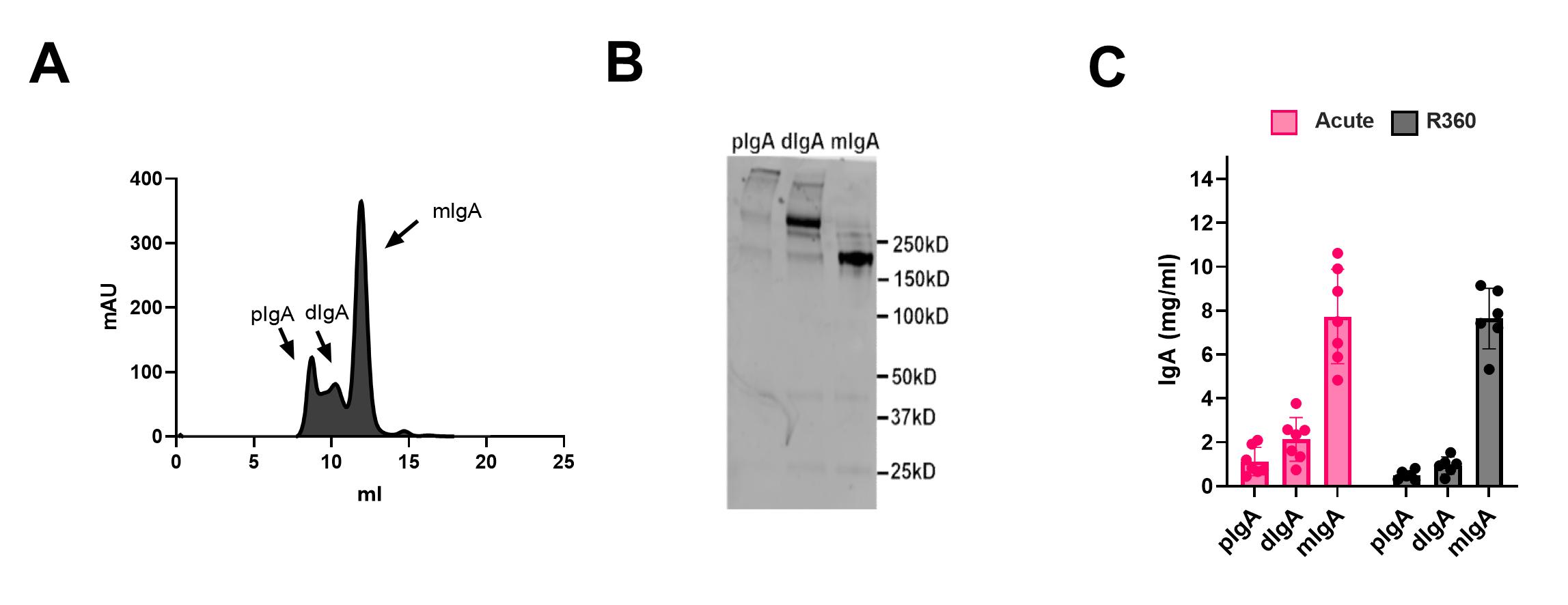

### Suppl Fig.5

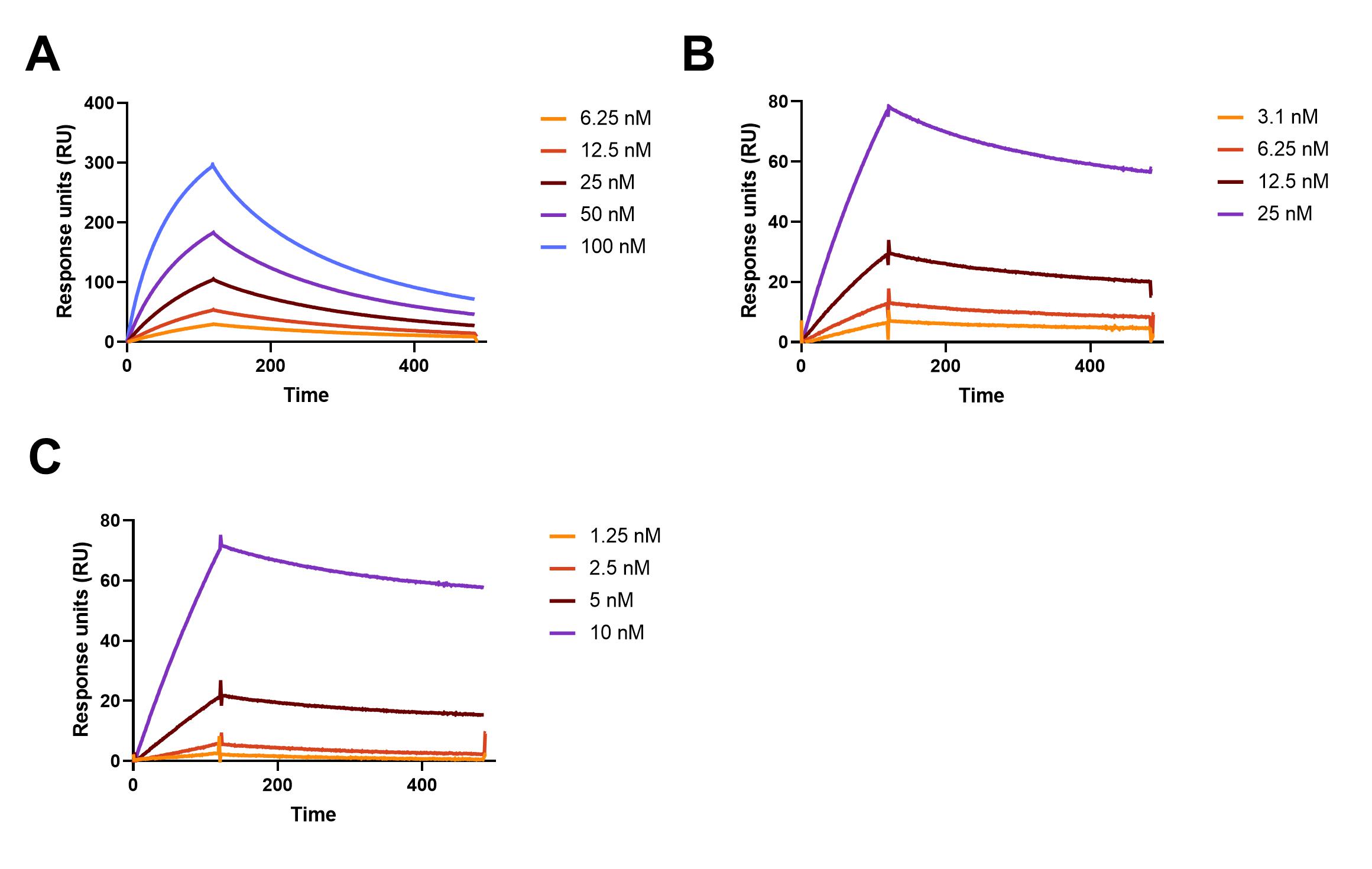

### Suppl Fig.6

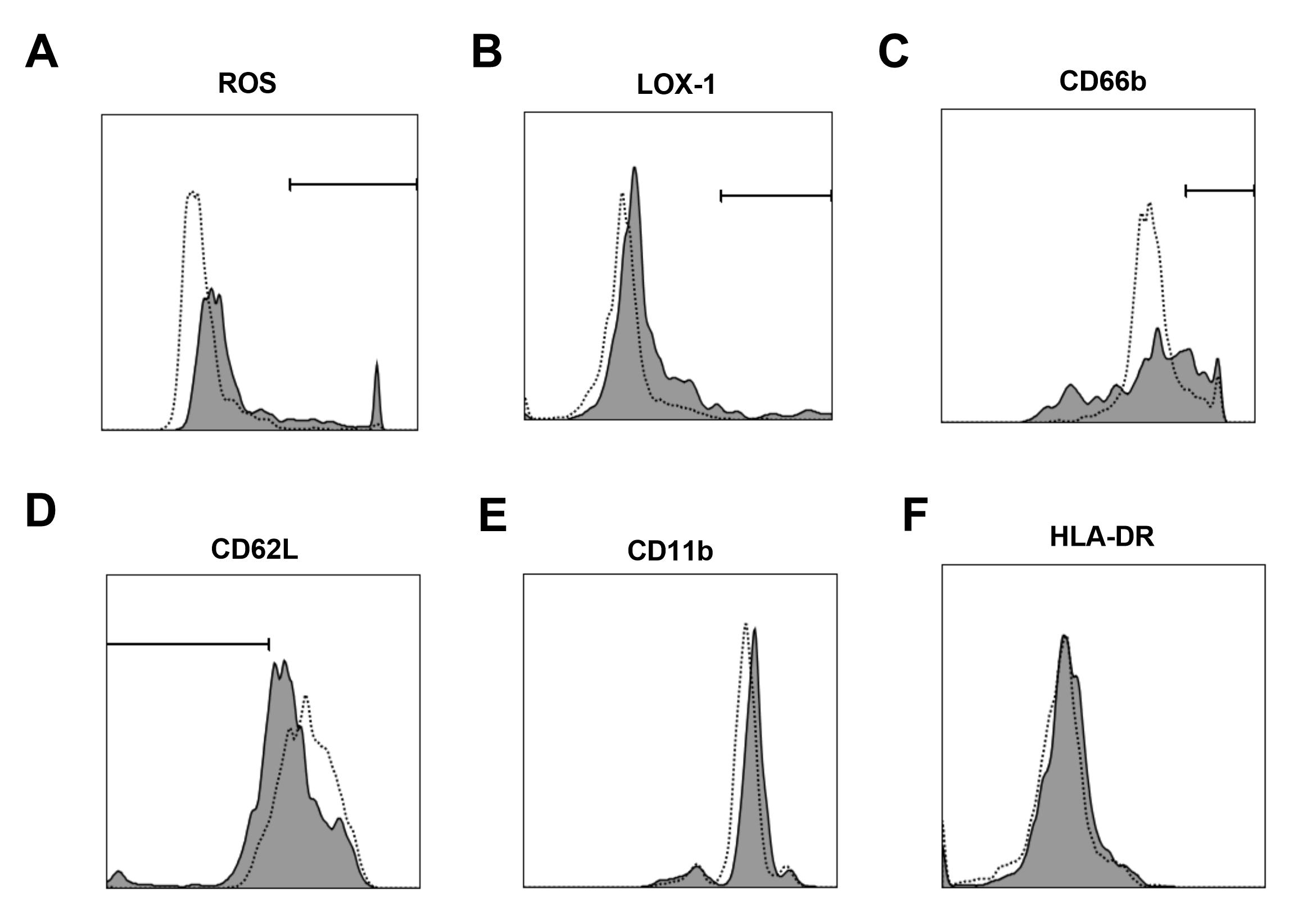
